# Disentangling Brain-Psychopathology Associations: A Systematic Evaluation of Transdiagnostic Latent Factor Models

**DOI:** 10.64898/2025.12.21.695029

**Authors:** Martin Gell, Mauricio S. Hoffmann, Tyler M. Moore, Aki Nikolaidis, Ruben C. Gur, Giovanni A. Salum, Michael P. Milham, Robert Langner, Veronika I. Müller, Simon B. Eickhoff, Theodore D. Satterthwaite, Brenden Tervo-Clemmens

## Abstract

Identifying robust brain-psychopathology associations with neuroimaging remains difficult, in part due to substantial heterogeneity within and comorbidity between diagnostic categories. Transdiagnostic latent factor models aim to address this structure by separating shared and unique symptom variance. However, it remains unclear whether latent factor modeling translates into stronger and more interpretable brain-psychopathology associations within contemporary whole-brain prediction frameworks. Using two large developmental cohorts, we systematically compared transdiagnostic bifactor models, correlated factor models, and typical summary scores derived from the Child Behaviour Checklist (CBCL) in their reliability and multivariate associations with whole-brain structure (MRI) and function (resting-state fMRI). General psychopathology, internalising, externalising, and attention dimensions could be significantly predicted from resting-state connectivity but not cortical thickness. We found no consistent evidence that latent factors (bifactor or correlated factor models) strengthened reliability or brain-psychopathology associations, relative to summary scores. Neural signatures were also highly consistent across all scoring methods, with triple network (DMN, FPN, CO) involvement in general, internalizing, and externalising psychopathology. Bifactor scores did, however, display more distinct neural signatures between general, internalising, and externalising dimensions than did summary or correlated factor scores. Collectively, results suggest that additional phenotypic modelling of psychopathology alone does not systematically strengthen the predictive utility of neuroimaging, possibly reflecting fundamental limits on the amount of explainable symptom variance by brain features. While latent factor models may aid in distinguishing neural correlates between constructs, improving phenotypic assessment may be necessary for improvements to brain-psychopathology association strength.

## Introduction

Widespread symptom comorbidity and diagnostic heterogeneity have long been recognised as major limitations of traditional, categorical psychiatric diagnoses (L. A. Clark et al., 1995; Newman et al., 1998). Comorbidity and heterogeneity also pose challenges for psychiatric neuroimaging studies that seek to link brain and psychopathology (Feczko & Fair, 2020). These challenges have motivated proposals to better capture psychopathology as latent dimensions with a hierarchical structure (Krueger, 1999; Neale & Kendler, 1995). Building on this work, comprehensive dimensional models like the Hierarchical Taxonomy of Psychopathology (HiTOP; Kotov et al., 2017) formalise psychopathology as a hierarchy of empirically derived spectra ranging from narrow symptom clusters to broad dimensions and a higher-order general factor.

Multiple statistical parameterisations can instantiate such hierarchical structures, including correlated factor models, higher-order factor models, and bifactor models (Reise, 2012; Kotov et al., 2017). Among these, bifactor models have been particularly prominent because they explicitly partition variance into a higher-order general “P” factor representing the general manifestation of psychopathology and orthogonal specific factors that can disentangle transdiagnostic and domain-specific processes (Lahey et al., 2012; Caspi et al., 2014). While not without criticism (c.f., Watts et al., 2020, 2024), bifactor models have been widely used to examine how common and unique features across symptoms relate to external outcomes like academic performance or cognition (Caspi & Moffitt, 2018; Smith et al., 2020). Importantly, recent perspectives (Zald & Lahey, 2017; Tiego et al., 2023) have proposed latent variable models, particularly bifactor models, may help address widespread challenges in identifying robust and reproducible brain-psychopathology relationships.

Factor analytic models offer potential advantages for studies linking brain and mental health phenotypes by improving the internal consistency, precision, and interpretability of measured constructs. These models extract latent dimensions of shared variance across symptoms, while accounting for unique variance and measurement error; together, these properties may lead to larger brain-psychopathology associations and enhance statistical power (Tiego et al., 2023). In contrast, typical, simple summary scores may inadvertently combine general, domain-level, and symptom-specific variance, producing heterogeneous phenotypes that obscure or dilute associations (Reise, 2012). This is critical because measurement imprecision and poor reliability attenuate brain-behaviour effect sizes and reduce the likelihood of detecting meaningful associations (Karvelis et al., 2023; Gell et al., 2024). Conversely, increasing reliability, while holding other parameters constant, may lead to increased signal-to-noise ratios, in turn improving brain-behaviour associations (Milham et al., 2021; Nikolaidis et al., 2022). Finally, compared to alternative correlated factor models (Kotov et al., 2017; Sunderland et al., 2021), bifactor approaches partition variance explicitly into uncorrelated general and specific factors (which may otherwise result in correlated factors). Such orthogonal factors can potentially reduce confounding of brain-psychopathology associations and help clarify whether brain correlates reflect general psychopathology or domain-specific liabilities (Zald & Lahey, 2017; Lahey et al., 2021). Nevertheless, the extent to which such psychometric advantages of bifactor models translate to stronger or more specific brain-psychopathology associations remains unexplored.

Prior studies have reported associations between bifactor-derived psychopathology dimensions and neuroimaging measures (Elliott et al., 2018; Kaczkurkin et al., 2018, 2019). However, because none have directly compared bifactor scores to alternative scoring approaches (e.g., summary scores), it remains unclear whether the identified brain correlates reflect novel insights into brain-psychopathology associations. Further, due to the large sample sizes required to detect robust and reproducible brain-behaviour associations (Marek et al., 2022), comprehensive evaluations of the relative utility of factor scores for brain-psychopathology modelling were, until recent large-scale consortia datasets, not feasible. Bifactor models themselves likewise require substantial sample sizes to fit, often resulting in factor scores being computed on the same data used to test for associations with brain imaging, introducing the risk of embedded circularity (“double dipping”), train-to-test leakage, and overfitting. There is also a lack of clarity in how best to incorporate latent factor models, typically estimated with structural equation modeling (SEM), into whole-brain, high-dimensional regularised prediction pipelines now considered best practice for reproducible brain-behaviour studies (Poldrack et al., 2020). Standard SEM frameworks are not designed to accommodate thousands of highly collinear imaging predictors used in whole-brain prediction. Studies therefore rely on estimated factor scores and report factor determinacy (FD) to quantify how well estimated scores approximate the true latent scores.

Although latent factor models have shown psychometric advantages in behavioural research, it remains unknown whether these advantages confer stronger or more distinct brain-psychopathology associations than typical summary scores. Here, we systematically evaluate the reliability and whole-brain multivariate predictive accuracy of factor scores from cortical thickness (obtained from structural MRI) and functional connectivity (obtained from resting-state fMRI). To evaluate the potential measurement benefits afforded by factor analysis, we benchmark prediction accuracy and multivariate feature weights of bifactor and correlated factor model scores against standard summary scores that do not require factor analysis (i.e., equally weighted sums of items). To ensure generalizability across model solutions, we utilised 11 previously bifactor and corresponding correlated factor models from item-level data of the Child Behavior Checklist (CBCL) in two large, diverse developmental samples: the Adolescent Brain Cognitive Development study (ABCD) (Volkow et al., 2018) and the Brazilian High-Risk Cohort (BHRC) (Salum et al., 2015).

## Methods

### Adolescent Brain Cognitive Development dataset

#### Participants

To investigate psychometric properties of latent factor models and their association with brain imaging, we used baseline and follow-up 1 data from the Adolescent Brain Cognitive Development study, a large longitudinal neuroimaging cohort study of 21 sites in the United States (Volkow et al., 2018). Only English-speaking participants without severe sensory, intellectual, medical, or neurological issues who completed all items of the CBCL at both time points (mean interval = 12.1 months) were selected. This resulted in a total of 10,897 participants (5698 female, ages = 9-11 at baseline) with complete CBCL data for both visits that were used to fit all factor models. A subset of ABCD participants who completed the baseline imaging session, finished all rs-fMRI sessions, and passed the ABCD quality control for their T1 and rs-fMRI were used for brain-psychopathology analyses. This subset comprised 6,572 participants (3,277 female, ages = 9-11).

#### Neuroimaging data, preprocessing and analyses

The ABCD MRI acquisition protocol (Casey et al., 2018) was harmonised across 21 sites on Siemens Prisma, Phillips, and GE 750 3T scanners. It included high-resolution T1w MRI images with a 32-channel head coil using a 3D MPRAGE sequence (TR = 2500 ms, 1.0 mm isotropic voxels). The rs-fMRI images were acquired using gradient-echo echo planar imaging (TR = 800 ms, 2.4 mm isotropic voxels) and included four 5-minute runs totalling 20 minutes.

Structural and functional data were pre-processed using the ABCD-BIDS pipeline, available through the ABCD-BIDS Community Collection (ABCC; Collection 3165) as detailed in Feczko et al. (2021). The preprocessing steps included distortion correction and alignment using Advanced Normalisation Tools (ANTS), FreeSurfer segmentation, and both surface and volumetric registration using FSL FLIRT rigid-body transformation. Resting-state fMRI data were further processed using the DCAN BOLD Processing (DBP) pipeline, which involved detrending, demeaning, and denoising via a general linear model incorporating tissue class and motion regressors. Following this, data were bandpass filtered between 0.008 and 0.09 Hz using a second-order Butterworth filter. Additional processing included respiratory motion filtering (targeting breathing rates between 18.58 and 25.73 breaths per minute) and censoring of frames exceeding a framewise displacement (FD) threshold of 0.2 mm or identified as statistical outliers (±3 standard deviations). The denoised time courses were parcellated using the HCP multimodal atlas with 360 cortical regions of interest (Glasser et al., 2016), together with 19 subcortical regions (Desikan et al., 2006). The signal time courses were averaged across all voxels of each parcel, and functional connectivity between them was calculated as Pearson correlation and Fisher Z-transformed. Region-wise cortical thickness was averaged across all vertices within each parcel of the HCP multimodal atlas.

### Brazilian High-Risk Cohort study dataset

To replicate our reliability analyses of factor and standard CBCL summary scores in a dataset with different characteristics, we used CBCL scores from 771 participants (334 female, ages = 6 - 14) from the Brazilian high-risk cohort study (Salum et al., 2015). All participants had completed all items of the Portuguese version of the CBCL at baseline and the first follow-up session (mean interval = 17 months). The BHRC is a school-based community cohort from the cities of São Paulo and Porto Alegre that is enriched with children with current symptoms and/or family history of psychiatric disorders (for details, see Salum et al., 2015).

### Common and Specific Variance of Psychopathology

#### Child Behavioural Checklist and summary score

The Child Behavioural Checklist (CBCL) (Achenbach, 1983) was used as the basis for calculating summary and factor scores reflecting various dimensions of psychopathology. The CBCL is a parent-reported assessment of 120 items/symptoms for subjects aged 6 to 18 using a 3-point scale (0 = not true; 1 = somewhat/sometimes true; 2 = very true/often). The CBCL organises scores into eight syndromic summary scores: anxious-depressed, withdrawn-depressed, somatic complaints, rule-breaking behaviour, aggressive behaviour, social problems, thought problems, and attention problems. Additionally, the scores can be combined into broader indices by summing up item-scores, such as internalising problems (comprising anxious-depressed, withdrawn-depressed, and somatic complaints) and externalising problems (comprising rule-breaking behaviour and aggressive behaviour) that have been informed by factor analysis. Finally, a total problems score comprises the linear, equally weighted sum of all items. Here, we utilised the T-score values for all aforementioned CBCL summary scores that are typically used in the literature and available with many datasets, including the ABCD. To evaluate the impact of item-level response rescoring used for factor score calculation (see below), we additionally calculated the linear sum of all summary scores using the rescored data. This was shown to have no impact on subsequent analyses (see Supplementary Figure 7). Two types of modelling approaches described below were applied to the item-level CBCL data; for a diagrammatic overview, see Supplementary Figure 1.

#### Bifactor models

Recent work has identified 11 different bifactor model solutions for the CBCL (Constantinou & Fonagy, 2019; Hoffmann et al., 2022). Therefore, to comprehensively estimate the test-retest reliability of factor scores, we have investigated all 11 reported models (Achenbach, 1983; Haltigan et al., 2018; McElroy et al., 2018; Deutz et al., 2020; Moore et al., 2020; D. A. Clark et al., 2021). To ensure model stability (Savalei, 2011; Finney & DiStefano, 2013; DiStefano et al., 2021), we first rescored items to only indicate the presence or absence of symptoms (i.e., somewhat/ sometimes true and very true/often were re-coded to both be 1), as the response frequency of “very often” was below 5% for 114 of 119 items which can lead to unstable numerical solutions. Within each bifactor model, all CBCL items present in the model definition (**Fig. 1**) were configured to load on a general “P-factor” (Caspi et al., 2014). Additionally, a subset of items was set to residually load on “specific factors” that depended on the given model (see Supplementary Table 1). Following typical bifactor approaches (Hoffmann et al., 2022), specific factors were not allowed to correlate with each other, nor with the general factor. Confirmatory factor analyses (CFA) were carried out using Mplus (Muthen & Muthen, 1998; Hallquist & Wiley, 2018) using delta parameterisation and weighted least squares with a diagonal weight matrix with standard errors and mean- and variance-adjusted chi-square test statistics (WLSMV) estimators. For fit indices, see Supplementary Tables 2 and 3. Factor scores were generated using a regression method, resulting in 11 P-factor scores and 38 specific factor scores per subject.

**Figure 1.**
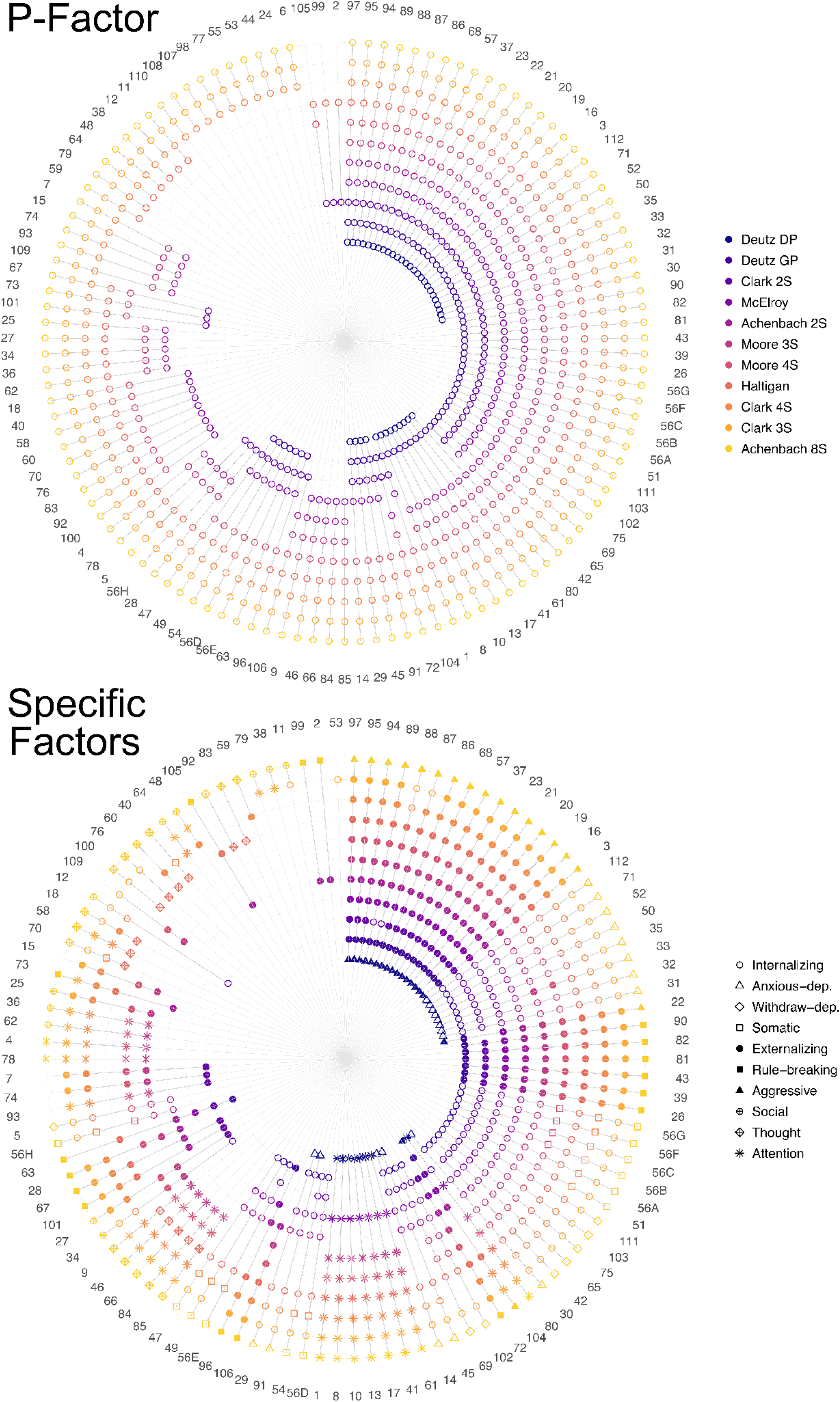
CBCL Items included in each model. Items included in the P-factor are depicted in the top panel, while specific factors are depicted in the bottom panel. Bifactor models included both P and specific factors, while correlated factor models only included specific factors. GP = general psychopathology model; DP = dysregulation profile model; CBCL = Child and Behaviour Checklist.

#### Correlated factor models

Correlated factor models were defined using the same data, 11 model solutions and CFA setup used for bifactor models. No general factor was defined, and specific factors were allowed to correlate. Due to near-perfect correlation with a number of other factors, the “social problems” factor was removed from the Achenbach 8S model. Therefore, in the ABCD, 37 specific factor scores per subject from 11 models were generated using a regression method. In the BHRC 2 models were removed due to non-convergence, resulting in 26 factor scores from 6 models.

### Reliability

We evaluated the test-retest reliability of factor scores across all models, as well as summed scores between the baseline and the first follow-up session for both the ABCD and BHRC samples. To this end, we calculated linear bivariate correlation, corrected for participant age, time point, and their interaction. Correlation may be more robust to systematic age-related changes in development, as it is not penalised by differences in means between baseline and follow-up data and different development rates across participants (Anokhin et al., 2022). Additionally, we calculated ICC using a two-way mixed-effects model for consistency, previously described as [3,1] (Shrout & Fleiss, 1979). In the ABCD, reliability was calculated on a subset of individuals who had a maximum of 12 months retest interval (n = 7250; 3774 female; retest interval mean = 11.3 months). To estimate reliability at shorter intervals, only subjects with a maximum of 6 months retest interval were selected from the BHRC sample, resulting in 234 subjects (100 female, retest interval mean = 3.7 months). To assess the internal consistency of factor scores, we calculated omega (ω), Hierarchical omega, and factor determinacy. For a detailed overview of each measure, see Supplementary Methods and Supplementary Table 4.

### Brain-psychopathology analyses

To systematically compare factor scores to CBCL summary T-scores with respect to their neurobiological substrates, we used functional connectivity and cortical thickness features in the ABCD sample to predict both types of scores. Predictions we performed using linear ridge regression implemented in the scikit-learn library (version 0.24.2), wrapped in custom code [https://github.com/MartinGell/Prediction_Psychopathology]. To avoid test-to-train leakage and improve generalizability, we utilised two matched samples (N = 3242 and 3330) created by Feczko et al. (2021) (so-called “discovery” and “replication” samples). These were matched on acquisition site, age, sex, ethnicity, grade, highest level of parental education, handedness, combined family income, and prior exposure to anaesthesia. Both bifactor and correlated factor models were fit separately within each sample to ensure factor score estimation remained independent across training and testing folds. Model evaluation was performed using a nested 2-fold cross-validation with 2 repeats, where each sample served once as training and once as testing data. Within each outer fold, the α regularisation parameter was optimised via efficient leave-one-out cross-validation (Rifkin & Lippert, 2007) on the training set, and performance was evaluated on the test fold. Sensitivity analyses using CBCL summary scores that did not require fitting separately on train and test sets showed that our 2-fold cross-validation yielded near-identical results to a more standard 5 times repeated 10-fold cross-validation (see Supplemental Methods for details).

Within each fold, neuroimaging features were z-scored across subjects (i.e. standard scaler) using training data, and the same transformation was applied to the test set using learned parameters from training data. To control for the effect of sex, given its common association with psychiatric phenotypes (Eaton et al., 2012), we performed feature-wise confound removal using linear regression (More et al., 2021). This was performed within each training fold, and the confound models were subsequently applied to test data to prevent data leakage. No other covariates were included due to the matched design. Prediction accuracy was quantified using Pearson correlation and the coefficient of determination (R²), which reflects explained variance and, within this framework, is not simply equivalent to the squared correlation coefficient (Poldrack et al., 2020). The significance of predictions was assessed using 1000 permutations of target labels. Feature weights (indicating which edges contributed more to predictions) were Haufe transformed (Haufe et al., 2014; J. Chen et al., 2023) to improve interpretability (see Supplementary Methods for details). To assess model sensitivity, we repeated all predictions using a gradient-boosted decision tree model (XGBoost) (T. Chen & Guestrin, 2016), which accommodates non-linearities and zero-inflated outcomes (see Supplemental Figure 2).

To examine the similarity between regression feature weights (i.e. neural correlates) between as well as within factor and summary scores, we correlated the upper triangles of the Haufe-transformed feature weight matrices. The significance of these similarities was evaluated using a cortex-only spin test (Alexander-Bloch et al., 2018; Markello & Misic, 2021), as the inclusion of subcortical parcels removes the possibility of surface projection. For each comparison, we computed the Spearman correlation (ρ) between the vectorised upper triangles of the two 360×360 matrices. Parcel centroids were projected to the spherical surface (fs_LR), and rigid-body rotations (“spins”) were applied while preserving left–right correspondence (Váša et al., 2018). For each spin, the resulting node permutation was applied to both rows and columns of one matrix, and ρ was recomputed. We repeated this procedure 10,000 times, each time recomputing ρ on the vectorised upper triangle to obtain the null distribution. To investigate the spatial embedding of the correlated connectomes, we used multidimensional scaling (see Supplementary Methods).

## Results

### Summary and factor scores show largely comparable test–retest reliability

Reliability and longitudinal stability were assessed using test-retest correlations corrected for participant age, time point, and their interaction. We compared the reliability of standard CBCL summary T-scores, which do not require factor analysis, to correlated and bifactor-derived factor scores from CBCL item-level responses based on 11 model solutions (see Methods). Reliability in the summary scores was higher in ABCD (mean across all scales: r_mean_ = 0.68, range: r = 0.56 - 0.76) than in the BHRC (r_mean_ = 0.53, r = 0.39 - 0.67), with total problems, externalising, and attention showing the greatest stability across both datasets. To compare corresponding constructs between summary and factor scores, we focus on the total summary score, P-factor, externalising, internalising and attention in the following sections (**Table 1**, for reliabilities of all scores see Supplementary Fig. 3 - 5, Supplementary Table 4).

**Table 1.** Test-retest correlation of corresponding constructs in summary and factor scores.

| Construct | ABCD |  |  | BHRC |  |  |
| --- | --- | --- | --- | --- | --- | --- |
|  | Summary score | Bifactor score mean (range) | Cor. factor score mean (range) | Summary score | Bifactor score mean (range) | Cor. factor score mean (range) |
| Total / P-factors | 0.76 | 0.74 (0.70-0.76) | -- | 0.60 | 0.58 (0.55-0.62) | -- |
| Externalising | 0.74 | 0.58 (0.54-0.62) | 0.74 (0.74-0.75) | 0.61 | 0.48 (0.40-0.54) | 0.49 (0.35-0.57) |
| Internalising | 0.68 | 0.57 (0.52-0.59) | 0.70 (0.68-0.71) | 0.53 | 0.37 (0.30-0.46) | 0.44 (0.30-0.60) |
| Attention | 0.74 | 0.56 (0.52-0.57) | 0.75 (0.72-0.76) | 0.67 | 0.45 (0.24-0.53) | 0.56 (0.49-0.62) |

As in published prior work, nearly all examined solutions for bifactor and correlated factor models generally had a good fit to the data in both ABCD and BHRC when considering multiple fit indices (Supplementary Table 2-3). Reliability was largely comparable between summary scores and corresponding constructs from bifactor and correlated factor models (**Table 1**) in both ABCD (n = 7250; retest interval mean = 11.3 months) and BHRC (n = 234; retest interval mean = 3.7 months and max = 6 months). Reliabilities were moderate to good for all constructs, except for the bifactor-derived specific factors such as externalising and internalising (ABCD: r = 0.52 - 0.62; BHRC: r = 0.24 - 0.54; see Supplementary Figs. 4 - 5). Factor determinacy was above 0.9 for all P-factors (FD_mea_ = 0.98, FD = 0.97 - 0.99) and most specific factors (FD_mean_ = 0.95, FD = 0.86 - 0.99) except for three factors from the Achenbach model with 8 factors (i.e., ACH8). Internal consistency reliability indices (ω, ωH) were high for P and low-to-acceptable for specific factors in both datasets (Supplementary Table 4). ICCs closely tracked test-retest correlations in both datasets and factor scoring methods (ABCD: r = 0.99, p < 0.001; BHRC: r = 0.99, p < 0.001).

### Summary and factor scores show comparable prediction accuracy

In the ABCD dataset, we used functional connectivity and cortical thickness fMRI/MRI features in a multivariate linear ridge regression to benchmark the prediction accuracy of brain-psychopathology associations. Overall, nearly all psychopathology constructs could be significantly predicted (assessed using permutation testing, see Methods) from functional connectivity (**Fig. 2A**, filled points). Prediction accuracies in line with previous work, explaining significant, but relatively low amounts of variance (see Supplemental Table 5), and were highly similar among summary (r_mean_ = 0.1, sd = 0.04; R²_mean_ = 0.01, sd = 0.007), bifactor (r_mean_ = 0.1, sd = 0.03; R²_mean_ = 0.008, sd = 0.008), and correlated factors (r_mean_ = 0.1, sd = 0.03; R²_mean_ = 0.01, sd = 0.007. For corresponding constructs (e.g., externalising factors vs externalising summary score), prediction accuracy for summary scores was likewise highly similar to the mean accuracy achieved for factor scores (**Fig. 2B**; for the coefficient of determination see Supplementary Fig. 6). The prediction accuracy of summary score t-scores typically used in the literature and raw sums across rescored items (see methods for details), used as the basis for factor score fitting, were near-identical (Supplementary Fig. 7).

**Figure 2.**
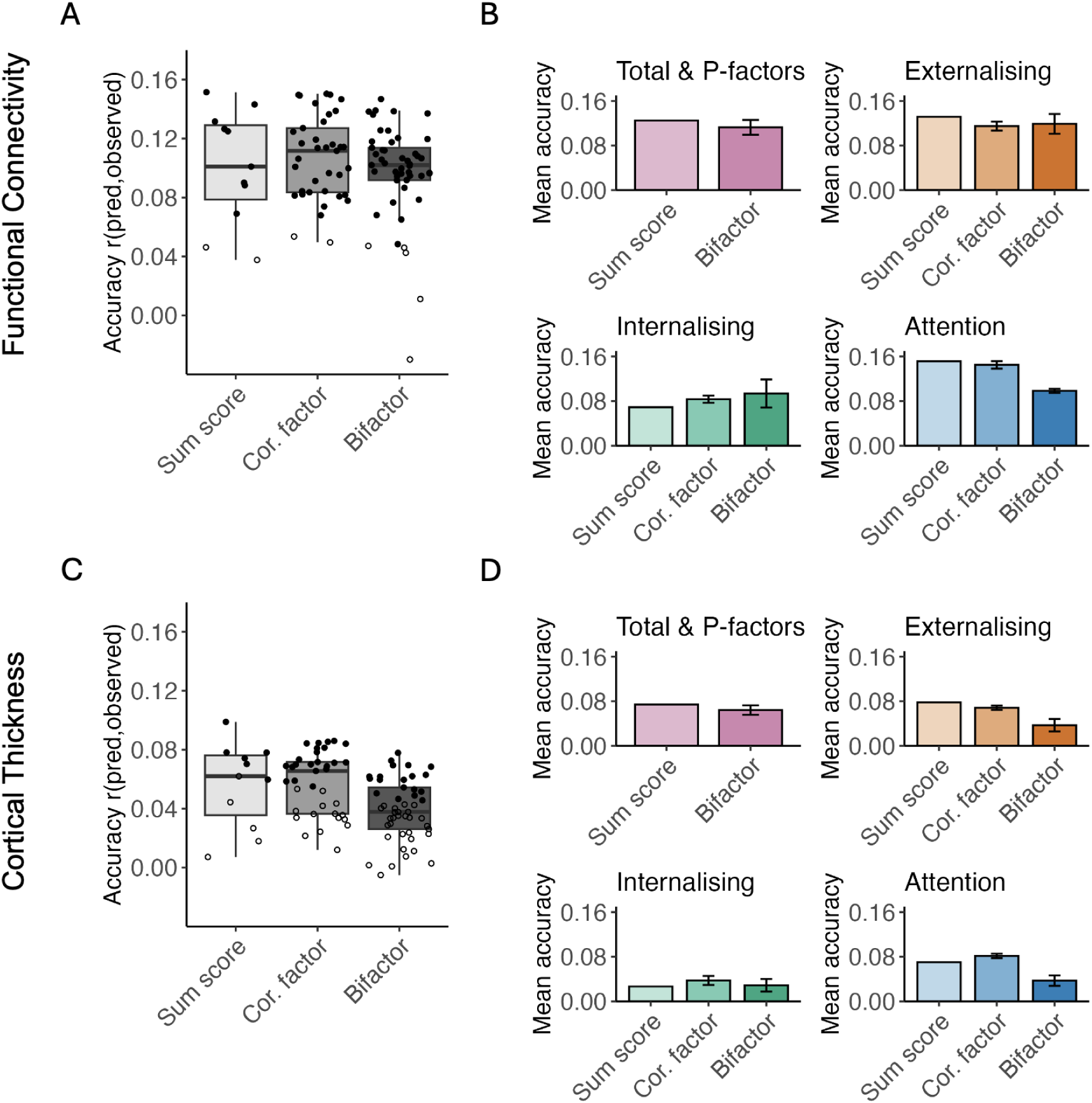
Prediction accuracy of CBCL summary and factor scores. The upper panel displays functional connectivity, and the lower panel shows cortical thickness-based prediction accuracy. Panels (A) and (C) show boxplots of prediction accuracies for all available summary scores, correlated factor and bifactor-derived factor scores (right). Panels (B) and (D) show the corresponding construct summary score and mean of correlated factor and bifactor score prediction accuracy. Error bars indicate standard deviation in accuracy across individual bifactor model solutions. Filled points in panels (A) and (C) represent permutation-based significant predictions at p < 0.001.

Prediction accuracy from cortical thickness was generally not significant (**Fig. 2C**, outline points) and produced near-chance results when evaluated using the coefficient of determination rather than correlation as a model performance metric (Supplementary Fig. 6). The majority of significant predictions were of the P-factors, which were also highly similar to the corresponding total problems (**Fig. 2D**; Supplementary Fig. 8; Supplementary Table 5). Owing to the overall poor predictive performance of cortical thickness, subsequent analyses focused on functional connectivity.

### Prediction accuracy scales with test-retest reliability, but not for the p-factor

Replicating prior work (Gell et al., 2024), summary scores with higher reliability had higher prediction accuracy (**Fig. 3A**). Similarly, within a given factor (i.e., P, externalising, internalising, attention), higher reliability also generally resulted in higher prediction accuracy across model solutions (**Fig. 3B-C**; each factor group illustrated by colour). However, for bifactor models, when comparing between factors (e.g., P vs. externalising), higher reliability did not necessarily translate to higher prediction accuracy. These results were consistent across predicted longitudinal timepoints in the ABCD dataset and machine learning algorithms (Supplementary Fig. 9).

**Figure 3.**
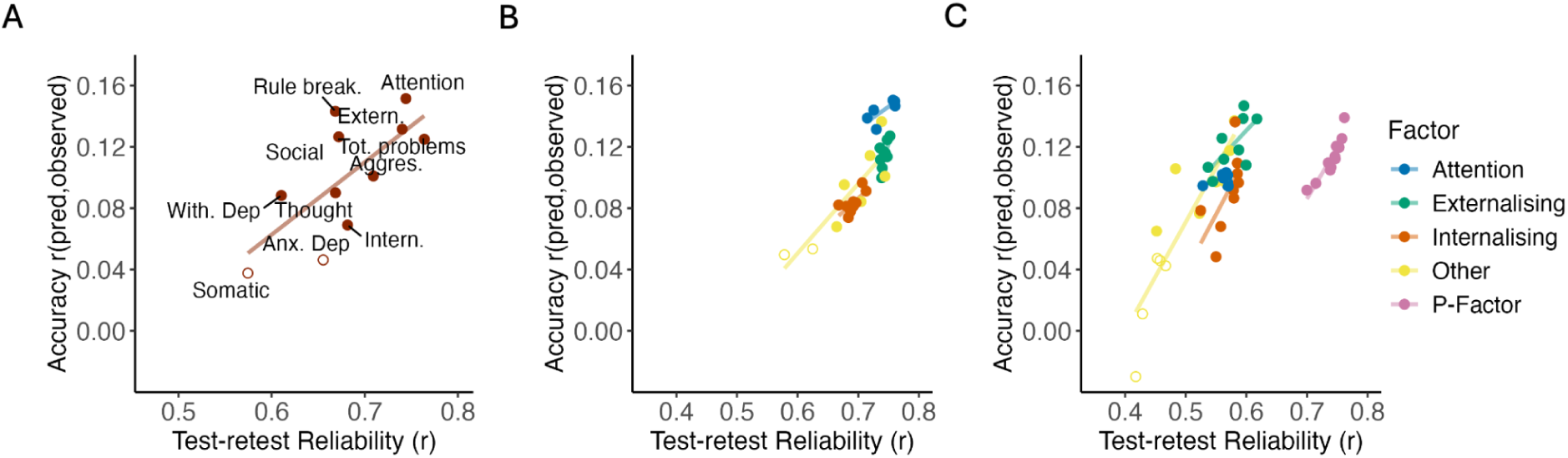
Impact of reliability on the prediction accuracy of factor scores from functional connectivity. The relationship between score test-retest reliability (x-axis) and prediction accuracy (y-axis; via linear ridge regression: see Methods) for summary scores (A), correlated factor scores (B) and bifactor scores (C). Filled points represent permutation-based significant predictions at p < 0.001. Each point refers to one model solution. For the impact of internal consistency reliability on prediction accuracy, see (Supplementary Fig. 11).

For bifactor models, despite having higher reliability and internal consistency than all other factors, P-factors could be predicted (r_mean_ = 0.11; R^2^_mean_ = 0.012) using functional connectivity with comparable accuracy to most specific factors (**Fig. 3C**; for the coefficient of determination see Supplementary Fig. 10). Externalising (r_mean_ = 0.12; R^2^_mean_ = 0.013) and attentional (r_mean_ = 0.10; R^2^_mean_ = 0.009) factors showed the most similar prediction strength to P-factors. Internalising displayed a slightly lower, yet still overlapping prediction accuracy (r_mean_ = 0.09; R^2^_mean_ = 0.002) to P-factors. Similarly to test-retest reliability, neither higher internal consistency reliability (ω, ωH, and factor determinacy) nor item variance explained by the corresponding factor could consistently index improvement to prediction accuracy for general compared to specific factors (Supplementary Fig. 11).

Collectively, these results underscore a distinction between and within constructs in neuroimaging-based prediction of bifactor models. One possibility is that there is a limited amount of overall predictable CBCL variance from brain imaging that places a theoretical ceiling on prediction accuracy. Importantly, this appears to be the case no matter how different bifactor models partition this variance into general or specific factors. Examination of all 11 bifactor models demonstrates differing proportions of item-level variance attributed to general and specific factors (**Fig. 4A**). Nevertheless, the average prediction accuracy from functional connectivity (**Fig. 4B**, grey line) across factors within each model solution was nearly identical (r = 0.10 - 0.12; R^2^ = 0.008 - 0.013).

**Figure 4.**
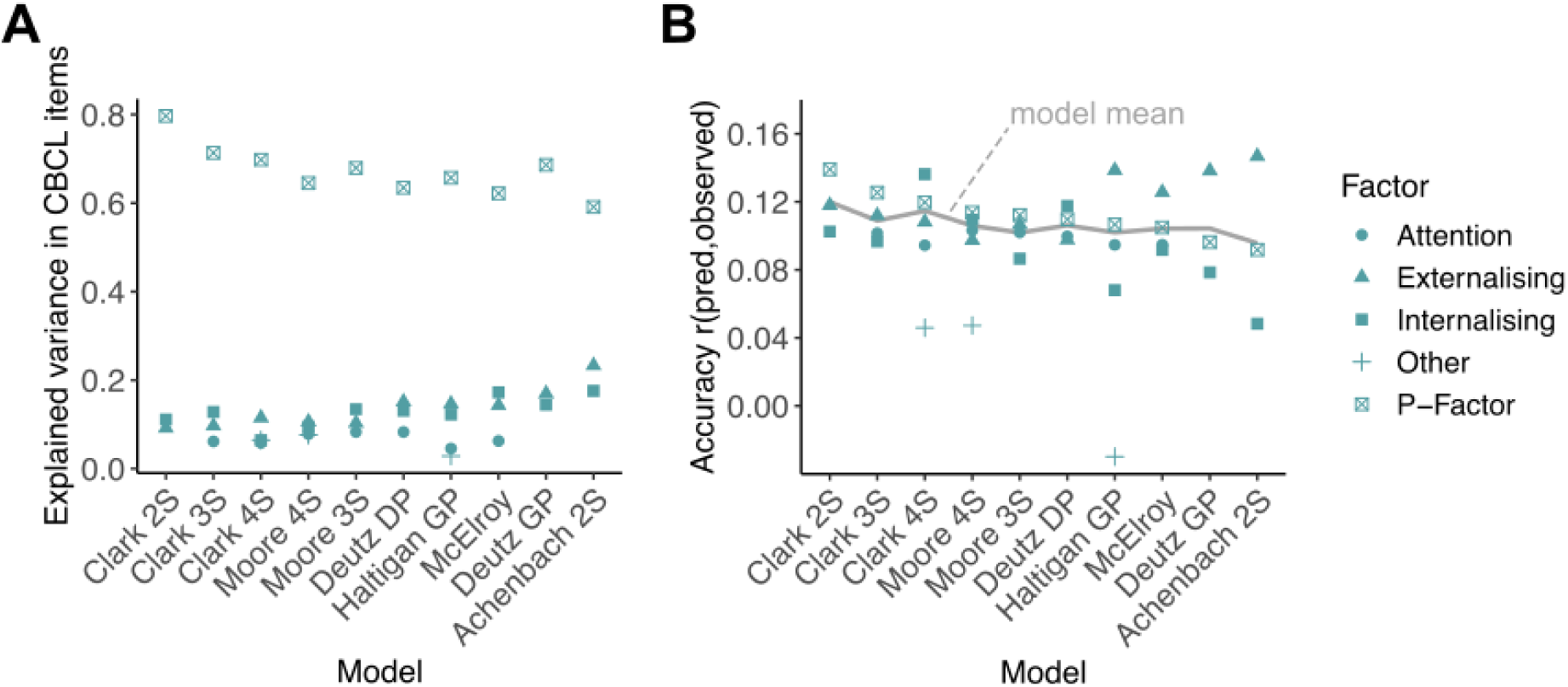
Explained item variance and prediction accuracy by functional connectivity. Results for whole-brain prediction of factor scores by linear ridge regression. Each point represents one factor. The Achenbach 8S model was removed as more than half of its specific factors could not be significantly predicted.

### Neural correlates of corresponding factor and summary score constructs show high similarity

Having shown that latent factor models did not enhance predictive performance relative to simple summary scores, we next tested whether these approaches might nevertheless reveal distinct neural correlates. To this end, we visualised the Haufe transformed feature weights (see supplementary methods for details) from our ridge regression prediction models, focusing on P-factors/total problems, externalising and internalising constructs (**Fig. 5** for summary and bifactor models; see Supplementary Figures 12 - 13 for comparison to correlated factor models). Connectivity within the default mode (DMN) as well as between the DMN and the frontoparietal (FPN) and cingulo-opercular (CO) networks were the most informative for predicting the total problems sum score (**Fig. 5A**). The externalising summary score prediction was most informed by an overlapping network of DMN and FPN edges with additional sensorimotor and attention network components. The prediction of the summary score of internalising symptoms was most informed by connectivity between visual, attention and FPN networks. This was also the case for the correlated factor model externalising and internalising.

**Figure 5.**
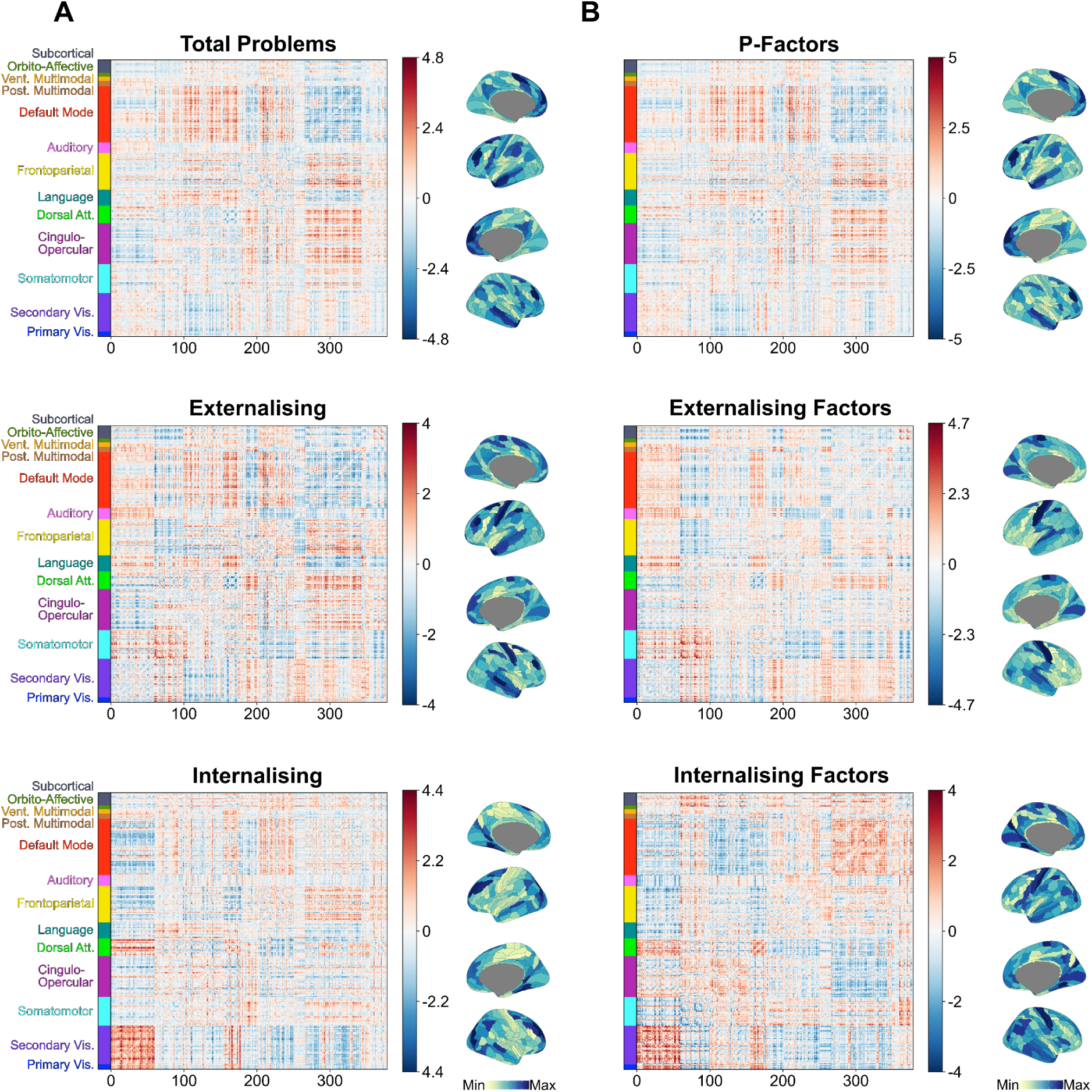
Haufe transformed feature importance weights of edges between all cortical parcels. Haufe-transformed feature weights for corresponding constructs of (A) summary score and (B) bifactor scores. In this case, positive or negative feature weight for an edge indicates that higher connectivity for that edge was associated with predicting higher or lower behavioural value, respectively. On the left side of each panel is the full feature weight matrix, ordered using the functional network definition by Ji et al. (2019). Brain maps display the mean absolute value weight for each cortical region. Note, weights for bifactor models are averaged across bifactor model solutions. For visualisation, the values within each matrix were divided by their standard deviations (across all entries in the matrix).

For latent factor models, feature weights across model solutions for a given construct (e.g. externalising) showed very high consistency (ρ_mean_ = 0.91 - 0.99; Supplementary Fig. 14 - 15) and were therefore averaged across models, resulting in one matrix of weights per construct. First, we compared the most informative features for the prediction of corresponding constructs from the summary and factor scores (**Fig. 6A**, highlighted diagonal values). This indicated a generally high similarity in feature weights between corresponding constructs (e.g., externalising factors vs externalising summary score). Functional connectivity features that predicted P-factors were almost perfectly spatially aligned with the total problems score (ρ = 0.98, p_spin_ < 0.001), also indicating DMN, FPN and CO network connections (**Fig. 5**). For bifactors, feature weights of externalising bifactors were likewise highly correlated with the externalising summary score (ρ = 0.85, p_spin_ < 0.001), mainly differing in the involvement of DMN connections in factor score predictions. Internalising bifactor and summary score feature weights showed the lowest, albeit still high similarity (ρ = 0.50, p_spin_ < 0.001), mostly differing in the increased importance of DMN and cingulo-opercular connectivity in factor score prediction. Correlated factors showed a near-perfect spatial similarity with feature weights of corresponding construct sum scores (externalising: ρ = 0.97, p_spin_ < 0.001; internalising: ρ = 0.94, p_spin_ < 0.001; Supplementary Figure 16).

**Figure 6.**
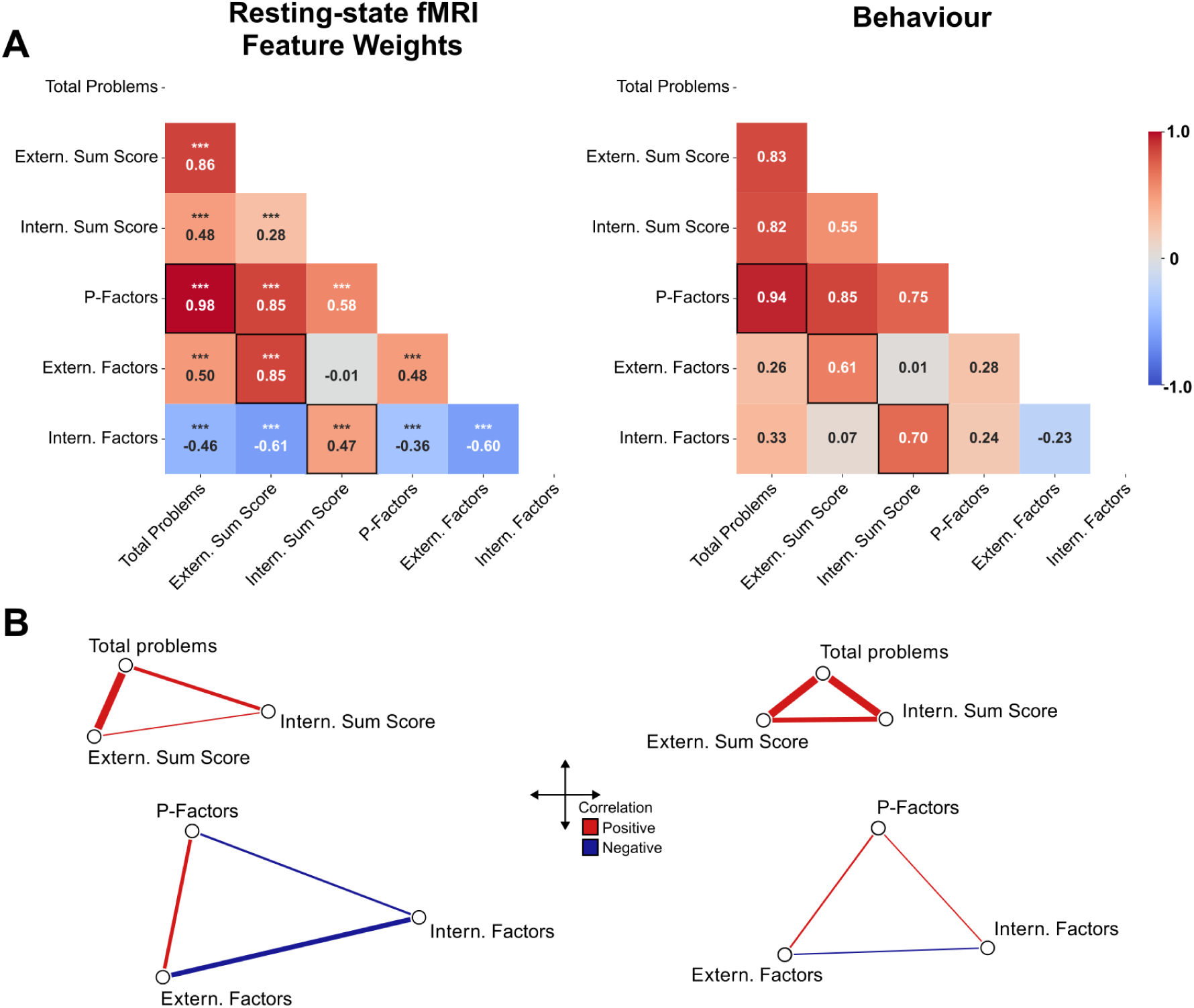
Correlation between informative features for the prediction of CBCL summary and factor scores. The left panel displays Spearman correlations between Haufe-transformed feature importance weights, while the right panel displays correlations between the actual phenotypic summary and factor scores used for prediction. Panel (A) shows the full matrix of between and within correlations. For any correlation involving the factor scores, the median across all correlations is displayed. Highlighted sections refer to correlations along the “theoretical diagonal”, i.e., between corresponding constructs (e.g., total summary and mean of P-factor weights). Significance p-values obtained using spin permutations. Panel (B) visualises the pattern of similarity (also shown as correlations in A) between general, externalising, and internalising constructs for summary (top) and factor (bottom) scores using a 2-D embedding computed from a distance matrix of correlations. Line thickness refers to correlation strength. See Supplementary Figures 16 and 17 for correlated factor models. Abbreviations: Extern: externalising; Intern: internalising; Prob: problems

The high similarity between corresponding factors was mirrored by high individual difference association between the factor and summary score phenotypes themselves (**Fig. 6A**; see Supplementary Fig. 18-20). The total problems score showed an almost perfect correlation with most bifactor-derived P-factors (ρ_mean_ = 0.94, ρ = 0.88 - 0.97). Externalising and internalising bifactors showed lower, albeit still high, correlations with the externalising (ρ_mean_ = 0.62; ρ = 0.49 - 0.69) and internalising (ρ_mean_ = 0.71; ρ = 0.57 - 0.75) summary scores, respectively. Correlated factors likewise showed near-perfect correlation with corresponding sum scores (externalising: ρ = 0.95; internalising: ρ = 0.93; Supplementary Figure 17).

### Neural correlates of bifactor scores display higher separability than correlated factor and summary scores

Finally, we investigated the feature weight similarity within each scoring approach (e.g., between bifactor-derived P-factor, externalising and internalising dimensions) to directly assess the cross-construct overlap and separability of neural correlates among psychopathology dimensions (**Fig. 6**). Correlation patterns among psychopathology constructs at the neural level closely tracked those at the individual differences of the phenotypes. Overall, at both the neural and phenotypic level, bifactor models had the lowest correlations between constructs, likely due to the orthogonalisation of factor scores. Outside of P-factor and externalising bifactor scores (ρ = 0.48, p_spin_ < 0.001), the most predictive features of bifactor scores showed negative correlations, indicating high dissimilarity in their neural correlates (**Fig. 6A**). In contrast, outside of internalising and externalising summary scores which showed a weak relationship between feature weights (ρ = 0.28, p_spin_ < 0.001) summary scores and correlated factor scores displayed high correlations on both the phenotypic and feature weight level (**Fig. 6A**; Supplementary Figures 16-17). Correlated factor score feature weights showed the highest similarity between externalising and internalising factors (ρ = 0.66, p_spin_ < 0.001). To visualise the pattern of similarity among factor and summary scores, we calculated a two-dimensional embedding of their similarity. These plots indicate that summary score feature weights and individual differences in phenotypic scores have higher similarity (i.e. have higher proximity in two-dimensional space) than factor scores (**Fig. 6B**). Overall, these results suggest that orthogonalisation of specific psychopathology dimensions may offer novel insights into neural correlates.

## Discussion

In this study, we investigated whether latent variable models of the CBCL confer measurable advantages for brain-psychopathology associations. Guided by the premise that latent factor models may strengthen or clarify brain-psychopathology associations by improving reliability and measurement precision, we compared scores from multiple latent factor models to simple summary scores. We found no consistent advantage of either bifactor or correlated factor models for the magnitude of brain-psychopathology prediction. Test-retest reliability was likewise comparable between correlated factor and summary scores from corresponding constructs, with the one exception that specific factors from the bifactor model (e.g., internalising, externalising, attention) had lower reliability than those from correlated factor models or summary scores. Feature weights from whole-brain predictive models of transdiagnostic psychopathology were broadly distributed across the connectome and consistent with prior theories emphasising higher-order networks (default mode, frontoparietal, and cingulo-opercular networks). However, these were likewise highly similar between factor scores and summary scores from corresponding constructs, with P-factors and total problems summary scores approaching numerical equivalence. One potential advantage of bifactor models over summary and correlated factor scores was that the pattern of neural correlates across constructs (e.g., p-factor vs. externalising vs. internalising) was more separable (i.e., less correlated), likely due to orthogonalisation of general and specific factors. This suggests bifactor scores in particular may provide novel insights (analogous to improved discriminant validity) into neural correlates, without substantial loss to prediction, though further work is necessary to adjudicate whether these insights are more or less valid.

The overarching results from this study challenge the assumption that factor analytic scores will inherently yield superior neurobiological insights, relative to simple summary scores. Prior work has proposed that hierarchical latent variable approaches like bifactor modelling can enhance reliability, interpretability and the robustness of brain-psychopathology associations by reducing measurement error and emphasising shared variance across symptoms, respectively (Zald & Lahey, 2017; Tiego et al., 2023). While the theory behind this is clear and may show practical gains in other contexts, the empirical pattern observed here suggests that factor models do not lead to systematically stronger predictions relative to simple summary scores. Rather, the small proportion of behavioural variance that can be explained by neuroimaging-derived brain features shown here and in the literature (J. Chen et al., 2022; Marek et al., 2022; Ooi et al., 2022; Heckner et al., 2023) may impose a ceiling on predictive accuracy irrespective of the scoring approach. In other words, psychometric refinements alone may not be sufficient to overcome fundamental constraints of effect size in large-scale brain-behaviour studies. Instead, a richer assessment of symptoms, environmental exposures, and developmental context (analogous to improving construct validity) may be necessary before reparameterization existing symptom inventories.

A remaining complexity in this work is how best to integrate latent factor models of psychopathology into whole-brain multivariate regularised prediction pipelines. Following previous work (Elliott et al., 2018; Kaczkurkin et al., 2018), our approach leveraged estimated factor scores, which are weighted composites of observed variables derived through regression or similar methods and are distinct from true latent factor scores, the unobservable values of the underlying construct (Rhemtulla & Savalei, 2025). Unlike latent factor scores, estimated factor scores carry residual indeterminacy (Grice, 2001) and do not fully eliminate measurement error, narrowing the expected gap between factor scores and simple summary scores. In the work here, the factor determinacy scores, which index how well estimated scores approximate their latent scores, were quite high, minimizing this concern. Nevertheless, future methodological research should rectify this issue of integrating latent models into brain-psychopathology prediction.

Our results indicate that greater reliability (test-retest, internal consistency and factor determinacy) for P-factors did not improve the strength of brain-behaviour associations compared to specific factors (e.g., externalising, internalising) with lower reliability. While these results suggest that general psychopathology symptoms are only weakly associated with brain structure and function, they also illustrate a fundamental distinction between reliability and construct validity: reliability is necessary but not sufficient for strong associations with external variables (Cronbach & Meehl, 1955). Psychopathology measures must index variance that is relevant to brain imaging (the external criterion), and increasing reliability does not necessarily increase this relevant variance. A similar principle can be illustrated for internal consistency reliability or FD: While general factors consistently demonstrated higher FD than specific factors, precision in estimating a latent construct did not guarantee better alignment with biologically meaningful variance. Nevertheless, reliability remains an important consideration for brain imaging of psychopathology – even if not sufficient, it is still necessary. For example, model solutions with higher reliability have better predictive performance than those with lower reliability for a given construct (e.g., across all P-factors), even if that does not generalise across constructs (i.e. P-factors vs. externalising factors).

The comparison of neural correlates of transdiagnostic psychopathology between summary and factor scores resulted in both overlapping and distinct network features underlying predictions. Most corresponding factor and summary score constructs share largely overlapping neural correlates, which aligns with prior evidence for broad ‘triple-network’ transdiagnostic connectivity patterns across youth psychopathology (Menon, 2011). Connectivity within and between the DMN, FPN and CO networks observed here has been consistently linked to mental health problems across samples (Xia et al., 2018; Sripada et al., 2021; Dhamala et al., 2023; Qu et al., 2023; Lee et al., 2024). Interestingly, while P-factors showed nearly identical feature weights and phenotypic scores with the total problems summary score (see also Fried et al., 2021), the bifactor-derived externalising and internalising scores were the only constructs where DMN-FPN interaction was not informative. Bifactor-derived Internalising scores were more reliant on DMN-CO connectivity, echoing work on the importance of the salience network and limbic regions in internalising symptoms (Menon, 2011; Cash et al., 2021; Pawlak et al., 2022). This divergence between bifactor and summary or correlated factor score findings is not surprising given the orthogonalisation of shared symptom variance from specific factors on the phenotypic level in the bifactor models. Furthermore, it highlights the potential benefits of higher separability in feature weights and behavioural data observed in bifactor models. However, it is also important to stress that while general and unique neural correlates may be informative, there is currently no ground truth to which they can be compared. Together, these findings suggest that while many constructs derived from the CBCL map onto a general, transdiagnostic network architecture, examining latent factors of specific symptom domains may reveal meaningful deviations in network topology.

Several limitations should be considered when interpreting these findings. First, all psychopathology measures were derived from parent-reported CBCL data, which may differentially capture externalising versus internalising behaviours. Externalising symptoms such as aggression or impulsivity are more readily observable, potentially inflating their predictive associations with neural features compared with less overt internalising symptoms (De Los Reyes & Kazdin, 2005; Rescorla et al., 2013). Second, our focus was on comparing factor and summary scores in their psychometric utility and association with brain imaging. Therefore, our findings of limited practical gains from factor models are specific to brain-psychopathology associations. It remains possible that factor scores could offer advantages over summary scores in other studies of criterion validity, such as predicting clinical outcomes or cognitive performance (e.g., McNeish, 2023). Alternatively, the lack of differential effects observed here may be driven by characteristics of the adolescent sample with relatively low base rates and limited severity of psychiatric symptoms. Such restricted individual variability may attenuate the ability to detect differences in brain-psychopathology associations between scoring approaches (Pavlovich et al., 2025; Gell et al., 2025). By extension, it remains plausible that stronger associations and more distinct patterns in neural correlates could emerge in contexts where psychopathology is more severe or prevalent, such as later developmental periods or in symptom-enriched cohorts (Kang et al., 2024).

Together, these findings suggest that latent factor models of psychopathology based on the CBCL offer limited added utility for explaining individual differences in brain structure and function beyond simple symptom summary scores. While latent factor modelling may improve psychometric precision and provide novel insights into neural correlates, its benefits for psychiatric neuroimaging appear constrained by inherently small effect sizes and are unlikely, on their own, to improve the prediction of mental health substantially. Improving phenotypic assessment, before exploring alternative phenotypic modelling, may provide more tangible improvements moving forward.

## Supporting information

Supplementary Methods and Results

