## Supplementary Methods and Results for "Disentangling Brain-Psychopathology Associations: A Systematic Evaluation of Transdiagnostic Latent Factor Models"

#### Supplement to Methods

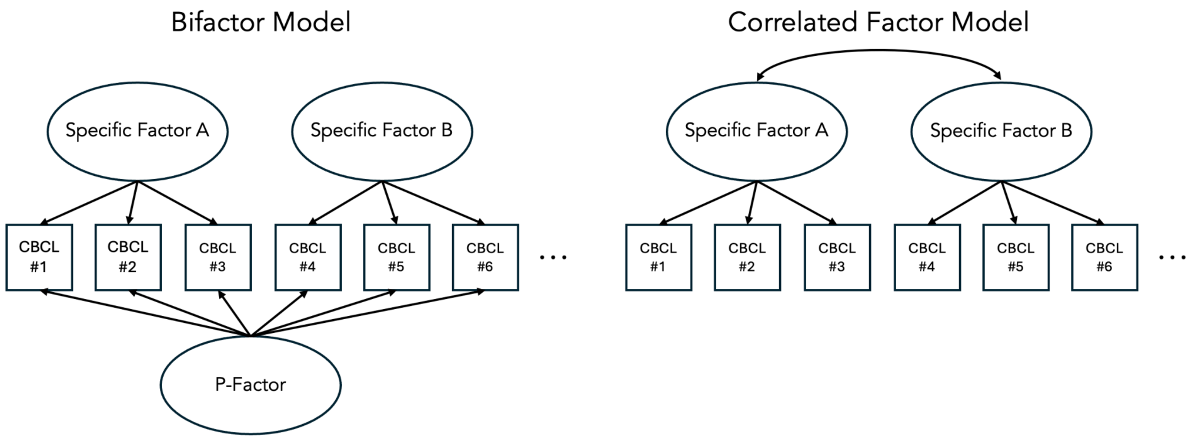


Supplementary Figure 1. Diagrammatic Model Overview. The left panel illustrates bifactor models, and the right panel illustrates correlated factor models.

##### Description of CBCL Bifactor Models

We evaluated 11 previously published CBCL bifactor model solutions (Supplementary Table 1) identified in prior systematic reviews and methodological syntheses (Constantinou & Fonagy, 2019; Hoffmann et al., 2022). Across solutions, items load on a general psychopathology factor ("P") and on one or more orthogonal specific factors (e.g., externalising, internalising, attention). Following prior work (Hoffmann et al., 2022), CBCL item responses were dichotomised to mitigate sparse high-severity response categories and to harmonise thresholds across items. Confirmatory factor analyses (CFA) used WLSMV estimation with delta parameterisation. Specific factors were constrained to be uncorrelated with one another and with the P-factor. Factor scores for all factors were exported for downstream reliability and prediction analyses. Correlated factor models were defined using the same data, 11 model solutions (Supplementary Table 1), and CFA setup used for bifactor models. No general factor was defined, and specific factors were allowed to correlate.

**Supplementary Table 1**

*Summary of CBCL Item-level Bifactor Models*

| Model | Factors | N items | Reference |
|---|---|---|---|
| Achenbach 2S bifactor model built upon ASEBA's 2 syndrome model | P-factor | 68 | Achenbach & Rescorla (2004) |
| | Internalizing | 33 | |
| | Externalizing | 35 | |
| Achenbach 8S bifactor model built upon ASEBA's 8 syndrome model | P-factor | 104 | Achenbach & Rescorla (2004) |
| | Anxious-depressed | 13 | |
| | Withdraw-depressed | 8 | |
| | Somatic | 12 | |
| | Rule breaking | 17 | |
| | Aggressive | 18 | |
| | Social problems | 11 | |
| | Thought problems | 15 | |
| | Attention problems | 10 | |
| Moore 3S | P-factor | 75 | Moore et al. (2020) |
| | Internalizing | 23 | |
| | Externalizing | 34 | |
| | Attention | 18 | |
| Moore 4S | P-factor | 75 | Moore et al. (2020) |
| | Internalizing | 16 | |
| | Externalizing | 34 | |
| | Attention | 18 | |
| | Somatic | 7 | |
| McElroy | P-factor | 66 | McElroy et al. (2018) |
| | Internalizing | 31 | |
| | Externalizing | 27 | |
| | Attention | 8 | |
| Deutz GP | P-factor | 71 | Deutz et al. (2020) |
| | Internalizing | 31 | |
| | Externalizing | 26 | |
| Deutz-Haltigan DP | P-factor | 39 | Deutz et al. (2020); Haltigan et al. (2018) |
| | Anxious-depressive | 13 | |
| | Aggressive | 18 | |
| | Attention | 8 | |
| Haltigan GP | P-factor | 85 | Haltigan et al. (2018) |
| | Internalizing | 31 | |
| | Externalizing | 32 | |
| | Thought problems | 14 | |
| | Attention | 8 | |
| Clark 2S | P-factor | 116 | Clark et al. (2021) |
| | Internalizing | 31 | |
| | Externalizing | 29 | |
| Clark 3S | P-factor | 116 | Clark et al. (2021) |
| | Internalizing | 43 | |

| | | | |
|---|---|---|---|
| | Externalizing | 32 | |
| | Attention | 25 | |
| Clark 4S | P-factor | 116 | Clark et al. (2021) |
| | Internalizing | 22 | |
| | Externalizing | 38 | |
| | Attention | 14 | |
| | Somatic | 12 | |

*CBCL = Child Behavior Checklist; S = bifactor model with n specific (S) factors;*

#### Cross-validation robustness

To assess whether our 2-fold cross-validation scheme with 2 repeats using so-called discovery and replication ARMS samples by Feczko et al. (2021), produced robust results, we benchmarked it against a standard 10-fold cross-validation repeated 5 times. This was only possible for the CBCL summary scores. As these are not model-based variables, there was no risk of leakage. Prediction accuracies were highly correlated across cross-validation schemes ($r(9) = 0.98$, $p < 0.001$). This suggests that the group-matched 2-fold design yielded results comparable to a more typical 10-fold cross-validation, while preserving a separation between the training and test folds.

#### Haufe transformation

Decoding models (e.g., ridge regression) produce weight vectors that are optimised for prediction but difficult to interpret as "importance" due to correlations among features. The Haufe transformation (Haufe et al., 2014) converts decoding weights into encoding-like maps that reflect each feature's association with the target while accounting for the covariance structure of the features. For a linear predictive model, the transformation is computed as the covariance of each feature and the predicted target variable in the training set. The transformed feature weights were averaged across folds to obtain the final weights. As we performed z-scoring of features as part of the ridge regression training in our main analyses, the training set features were also z-scored prior to computing the transformation. For more details on the Haufe transform, see (Haufe et al., 2014; J. Chen et al., 2023).

#### Extreme Gradient Boosting (XGBoost) analyses

To assess if the prediction of factor scores was impacted by the use of the linear ridge model, we repeated all predictions using a distributed gradient-boosted decision tree model that allows for non-linear relationships between features and the outcome. We used an extreme gradient boosting (XGBoost) regressor (T. Chen & Guestrin, 2016) to model the relationship between neuroimaging features and psychopathology factor/summary scores, as it is a non-linear model that is well-suited for handling zero-inflated distributions observed for some of the factor scores (Supplementary Figure 2). The XGBoost algorithm implemented in Python (available online at: https://github.com/dmlc/xgboost) is a tree-based ensemble method that iteratively builds decision trees while minimising a specified loss function (in our case, the mean squared error), incorporating second-order gradient information to improve convergence. Additionally, XGBoost implements shrinkage (i.e., learning rate) and column subsampling to improve predictive performance and computational efficiency. Specifically, we used random samples of 80% of the data for each tree, and limited the feature space for each split to 10% of the available features to reduce overfitting. Hyperparameters, including the learning rate, maximum tree depth, and number of estimators, were optimised using a nested, five-times-repeated, 3-fold cross-validation. The hyperparameter space and implementation are available in the GitHub repository containing all code used for analyses.

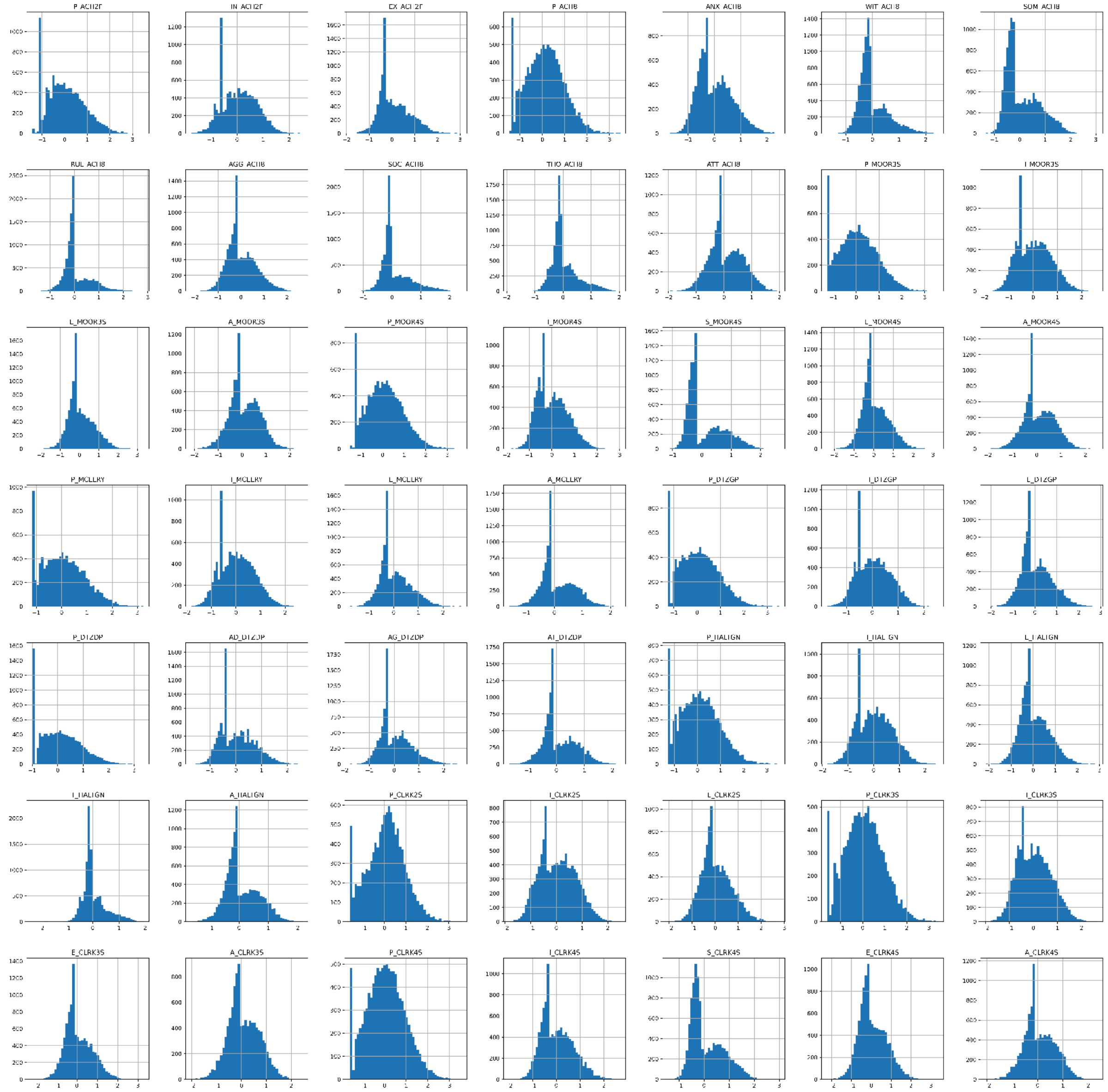

Supplementary Figure 2. Bifactor score distributions

#### 2-D embedding of correlations within summary and factor scores

The correlations within factor and summary scores were visualised as a network using custom R code [https://github.com/MartinGell/Prediction_Psychopathology]. Correlations were first converted to Euclidean distances. Next, each connected component of the resulting adjacency graph was embedded in two dimensions using multidimensional scaling (MDS) and centred without rescaling. Components were positioned side-by-side for visualisation.

### Supplement to Results

#### Factor score fit indices

Bifactor models had a generally good fit to the data in both datasets (Supplementary tables 2 and 3)

**Supplementary Table 2**

*Fit indices of all CBCL models for ABCD*

| Factor | Model | RMSEA | RMSEA 90% CI | CFI | TLI | SRMR |
|---|---|---|---|---|---|---|
| Bifactor | Achenbach 2S | 0.015 | [0.014, 0.015] | 0.994 | 0.993 | 0.088 |
| | Achenbach 8S | 0.018 | [0.018, 0.019] | 0.987 | 0.987 | 0.091 |
| | Moore 3S | 0.020 | [0.020, 0.020] | 0.994 | 0.994 | 0.078 |
| | Moore 4S | 0.020 | [0.020, 0.020] | 0.994 | 0.994 | 0.078 |
| | McElroy | 0.018 | [0.018, 0.019] | 0.994 | 0.994 | 0.078 |
| | Deutz GP | 0.017 | [0.016, 0.017] | 0.994 | 0.994 | 0.080 |
| | Deutz Haltigan DP | 0.027 | [0.027, 0.028] | 0.998 | 0.998 | 0.070 |
| | Haltigan GP | 0.016 | [0.016, 0.016] | 0.992 | 0.992 | 0.085 |
| | Clark 2S | 0.021 | [0.021, 0.021] | 0.981 | 0.980 | 0.089 |
| | Clark 3S | 0.015 | [0.015, 0.015] | 0.991 | 0.991 | 0.073 |
| | Clark 4S | 0.016 | [0.015, 0.016] | 0.990 | 0.989 | 0.070 |
| Correlated Factors | Achenbach 2S | 0.025 | [0.025, 0.025] | 0.981 | 0.981 | 0.105 |
| | Achenbach 8S | 0.102 | [0.102, 0.103] | 0.741 | 0.735 | 0.183 |
| | Moore 3S | 0.029 | [0.029, 0.029] | 0.987 | 0.987 | 0.086 |
| | Moore 4S | 0.029 | [0.029, 0.029] | 0.988 | 0.988 | 0.083 |
| | McElroy | 0.029 | [0.029, 0.029] | 0.984 | 0.984 | 0.092 |
| | Deutz GP | 0.134 | [0.134, 0.134] | 0.633 | 0.624 | 0.210 |
| | Deutz Haltigan DP | 0.048 | [0.047, 0.048] | 0.993 | 0.993 | 0.101 |
| | Haltigan GP | 0.021 | [0.021, 0.021] | 0.986 | 0.986 | 0.095 |
| | Clark 2S | 0.133 | [0.132, 0.133] | 0.240 | 0.233 | 0.282 |
| | Clark 3S | 0.058 | [0.057, 0.058] | 0.857 | 0.855 | 0.141 |
| | Clark 4S | 0.071 | [0.071, 0.072] | 0.781 | 0.777 | 0.168 |

CBCL = Child Behavior Checklist; S = specific factors; GP = general psychopathology; DP = dysregulation profile; RMSEA = root mean square error of approximation; CFI = comparative fit index; TLI = Tucker-Lewis index; SRMR = standarized root mean-square residual.

**Supplementary Table 3**

*Fit indices of all CBCL models for BHRC*

| Factor | Model | RMSEA | RMSEA 90% CI | CFI | TLI | SRMR |
|---|---|---|---|---|---|---|
| Bifactor | Achenbach 2S | 0.016 | [0.015, 0.017] | 0.993 | 0.992 | 0.103 |
| | Achenbach 8S | 0.021 | [0.020, 0.021] | 0.979 | 0.979 | 0.105 |
| | Moore 3S | 0.016 | [0.014, 0.017] | 0.994 | 0.994 | 0.095 |
| | Moore 4S | 0.017 | [0.015, 0.018] | 0.993 | 0.993 | 0.096 |
| | McElroy | 0.014 | [0.013, 0.016] | 0.993 | 0.993 | 0.092 |
| | Deutz GP | 0.017 | [0.016, 0.018] | 0.992 | 0.991 | 0.094 |
| | Deutz Haltigan DP | 0.017 | [0.015, 0.019] | 0.995 | 0.995 | 0.088 |
| | Haltigan GP | 0.017 | [0.016, 0.018] | 0.989 | 0.988 | 0.100 |
| | Clark 2S | 0.020 | [0.019, 0.021] | 0.977 | 0.976 | 0.103 |
| | Clark 3S | 0.018 | [0.018, 0.019] | 0.981 | 0.980 | 0.097 |
| | Clark 4S | 0.020 | [0.019, 0.020] | 0.978 | 0.977 | 0.100 |
| Correlated Factors | Achenbach 2S | 0.025 | [0.024, 0.026] | 0.981 | 0.980 | 0.118 |
| | Achenbach 8S | -- | -- | -- | -- | -- |
| | Moore 3S | 0.021 | [0.020, 0.022] | 0.988 | 0.988 | 0.102 |
| | Moore 4S | 0.021 | [0.020, 0.022] | 0.989 | 0.988 | 0.100 |
| | McElroy | 0.022 | [0.021, 0.024] | 0.983 | 0.983 | 0.106 |
| | Deutz GP | 0.117 | [0.116, 0.118] | 0.603 | 0.594 | 0.221 |
| | Deutz Haltigan DP | 0.023 | [0.020, 0.025] | 0.992 | 0.991 | 0.103 |
| | Haltigan GP | -- | -- | -- | -- | -- |
| | Clark 2S | 0.112 | [0.111, 0.112] | 0.258 | 0.251 | 0.276 |
| | Clark 3S | 0.056 | [0.055, 0.056] | 0.819 | 0.816 | 0.149 |
| | Clark 4S | 0.061 | [0.060, 0.061] | 0.782 | 0.779 | 0.174 |

CBCL = Child Behavior Checklist; S = specific factors; GP = general psychopathology; DP = dysregulation profile; RMSEA = root mean square error of approximation; CFI = comparative fit index; TLI = Tucker-Lewis index; SRMR = standarized root mean-square residual.

#### Factor internal consistency indices

To assess the internal consistency of factor scores in the ABCD, we calculated omega (ω), Hierarchical omega, and factor determinacy (Supplementary Table 4). Omega is a model-based analogue to the alpha coefficient, adjusting for varying factor loadings, and represents the associations between a test's items and its target construct. Similarly, Hierarchical omega (Hω) captures the proportion of total variance attributed to the general or specific factors. Generally, Hω > 0.8 is considered sufficient for interpretation (Rodriguez et al., 2016). Finally, factor determinacy (FD) assesses the correlation between the factor scores and the estimated factor and thus represents the extent to which factor scores are a good estimate of individual differences on the factor (Grice, 2001). Conventionally, FD > 0.9 is required for factor scores to be used (Gorsuch, 1980). FD was acceptable for P-factors and specific factors from all model solutions ($FD_{mean}$ = 0.98, FD = 0.96 - 0.99), except for three specific factors (0.86 - 0.87) from the Achenbach model with 8 factors (i.e., ACH8).

**Supplementary Table 4**

*CBCL bifactor level indices for all models in the ABCD*

| Model | Factor | r* | ICC(3,1) | ICC 95% CI | Omega | Omega H | FD | ECV_GS | ECV_SG | ECV_SS |
|---|---|---|---|---|---|---|---|---|---|---|
| Achenbach 2S | P-factor | 0.694 | 0.694 | [0.684, 0.703] | 0.931 | 0.685 | 0.975 | 0.591 | 0.591 | 0.591 |
| | Internalising | 0.543 | 0.544 | [0.530, 0.557] | 0.880 | 0.357 | 0.951 | 0.585 | 0.176 | 0.415 |
| | Externalising | 0.594 | 0.594 | [0.578, 0.605] | 0.915 | 0.283 | 0.969 | 0.596 | 0.233 | 0.404 |
| Achenbach 8S | P-factor | 0.745 | 0.745 | [0.737, 0.753] | 0.956 | 0.889 | 0.989 | 0.653 | 0.653 | 0.653 |
| | Anxious-depressed | 0.575 | 0.575 | [0.563, 0.588] | 0.827 | 0.313 | 0.942 | 0.609 | 0.051 | 0.391 |
| | Withdraw-depressed | 0.526 | 0.526 | [0.509, 0.539] | 0.724 | 0.251 | 0.868 | 0.621 | 0.028 | 0.379 |
| | Somatic | 0.446 | 0.446 | [0.431, 0.461] | 0.73 | 0.467 | 0.977 | 0.355 | 0.062 | 0.645 |
| | Rule Breaking | 0.482 | 0.482 | [0.467, 0.496] | 0.843 | 0.222 | 0.979 | 0.664 | 0.056 | 0.336 |
| | Aggressive | 0.540 | 0.540 | [0.525, 0.553] | 0.885 | 0.200 | 0.926 | 0.721 | 0.054 | 0.279 |
| | Social problems | 0.452 | 0.452 | [0.436, 0.466] | 0.758 | 0.104 | 0.927 | 0.752 | 0.025 | 0.248 |
| | Thought problems | 0.422 | 0.421 | [0.405, 0.436] | 0.77 | 0.045 | 0.855 | 0.823 | 0.021 | 0.177 |
| | Attention problems | 0.557 | 0.557 | [0.543, 0.569] | 0.846 | 0.258 | 0.979 | 0.589 | 0.050 | 0.411 |
| Moore 3S | P-factor | 0.742 | 0.742 | [0.733, 0.750] | 0.949 | 0.824 | 0.984 | 0.679 | 0.679 | 0.679 |
| | Internalising | 0.575 | 0.576 | [0.562, 0.588] | 0.848 | 0.432 | 0.949 | 0.471 | 0.134 | 0.529 |
| | Externalising | 0.535 | 0.535 | [0.520, 0.547] | 0.928 | 0.131 | 0.934 | 0.787 | 0.103 | 0.213 |
| | Attention | 0.553 | 0.553 | [0.539, 0.565] | 0.874 | 0.187 | 0.991 | 0.681 | 0.083 | 0.319 |
| Moore 4S | P-factor | 0.741 | 0.741 | [0.732, 0.749] | 0.951 | 0.831 | 0.982 | 0.645 | 0.645 | 0.645 |
| | Internalising | 0.583 | 0.584 | [0.569, 0.595] | 0.835 | 0.397 | 0.935 | 0.515 | 0.095 | 0.485 |
| | Somatic | 0.430 | 0.431 | [0.415, 0.446] | 0.788 | 0.584 | 0.993 | 0.214 | 0.076 | 0.786 |
| | Externalising | 0.542 | 0.542 | [0.528, 0.555] | 0.929 | 0.152 | 0.936 | 0.771 | 0.106 | 0.229 |
| | Attention | 0.557 | 0.557 | [0.544, 0.570] | 0.873 | 0.181 | 0.982 | 0.684 | 0.078 | 0.316 |
| McElroy | P-factor | 0.730 | 0.730 | [0.721, 0.739] | 0.936 | 0.750 | 0.972 | 0.622 | 0.622 | 0.622 |
| | Internalising | 0.572 | 0.573 | [0.560, 0.585] | 0.874 | 0.380 | 0.955 | 0.557 | 0.173 | 0.443 |
| | Externalising | 0.557 | 0.557 | [0.542, 0.569] | 0.913 | 0.247 | 0.930 | 0.687 | 0.143 | 0.313 |
| | Attention | 0.557 | 0.557 | [0.542, 0.569] | 0.799 | 0.272 | 0.929 | 0.594 | 0.063 | 0.406 |
| Deutz GP | P-factor | 0.710 | 0.711 | [0.701, 0.720] | 0.933 | 0.770 | 0.979 | 0.686 | 0.686 | 0.686 |
| | Internalising | 0.516 | 0.517 | [0.503, 0.530] | 0.875 | 0.278 | 0.946 | 0.641 | 0.144 | 0.359 |
| | Externalising | 0.614 | 0.614 | [0.600, 0.625] | 0.908 | 0.299 | 0.945 | 0.611 | 0.169 | 0.389 |
| Deutz DP | P-factor | 0.726 | 0.726 | [0.717, 0.735] | 0.925 | 0.769 | 0.966 | 0.635 | 0.635 | 0.635 |
| | Anxious-depressed | 0.567 | 0.567 | [0.554, 0.580] | 0.826 | 0.345 | 0.946 | 0.573 | 0.131 | 0.427 |
| | Aggressive behaviour | 0.559 | 0.559 | [0.544, 0.571] | 0.888 | 0.228 | 0.914 | 0.679 | 0.151 | 0.321 |
| | Attention | 0.551 | 0.551 | [0.536, 0.563] | 0.797 | 0.244 | 0.918 | 0.627 | 0.083 | 0.373 |
| Haltigan GP | P-factor | 0.734 | 0.734 | [0.726, 0.743] | 0.944 | 0.805 | 0.983 | 0.657 | 0.657 | 0.657 |
| | Internalising | 0.552 | 0.553 | [0.539, 0.565] | 0.874 | 0.324 | 0.958 | 0.602 | 0.122 | 0.398 |
| | Externalising | 0.591 | 0.591 | [0.576, 0.602] | 0.921 | 0.262 | 0.953 | 0.647 | 0.146 | 0.353 |
| | Thought | 0.408 | 0.407 | [0.391, 0.423] | 0.772 | 0.055 | 0.869 | 0.818 | 0.028 | 0.182 |
| | Attention | 0.517 | 0.517 | [0.500, 0.529] | 0.795 | 0.236 | 0.928 | 0.627 | 0.046 | 0.373 |
| Clark 2S | P-factor | 0.757 | 0.757 | [0.748, 0.764] | 0.955 | 0.888 | 0.992 | 0.797 | 0.797 | 0.797 |
| | Internalising | 0.579 | 0.580 | [0.565, 0.591] | 0.880 | 0.308 | 0.962 | 0.612 | 0.111 | 0.388 |
| | Externalising | 0.576 | 0.576 | [0.563, 0.588] | 0.903 | 0.197 | 0.944 | 0.702 | 0.092 | 0.298 |
| Clark 3S | P-factor | 0.752 | 0.752 | [0.744, 0.760] | 0.956 | 0.848 | 0.984 | 0.714 | 0.714 | 0.714 |

| | | | | | | | | | | |
|---|---|---|---|---|---|---|---|---|---|---|
| | Internalising | 0.580 | 0.580 | [0.566, 0.592] | 0.899 | 0.275 | 0.957 | 0.634 | 0.128 | 0.366 |
| | Externalising | 0.562 | 0.562 | [0.548, 0.574] | 0.918 | 0.196 | 0.944 | 0.718 | 0.097 | 0.282 |
| | Attention | 0.559 | 0.558 | [0.545, 0.571] | 0.872 | 0.092 | 0.972 | 0.753 | 0.061 | 0.247 |
| | P-factor | 0.748 | 0.748 | [0.740, 0.756] | 0.956 | 0.869 | 0.985 | 0.698 | 0.698 | 0.698 |
| | Internalising | 0.575 | 0.576 | [0.562, 0.588] | 0.863 | 0.248 | 0.942 | 0.684 | 0.064 | 0.316 |
| Clark 4S | Externalising | 0.435 | 0.435 | [0.420, 0.450] | 0.926 | 0.228 | 0.951 | 0.692 | 0.115 | 0.308 |
| | Somatic | 0.598 | 0.598 | [0.584, 0.609] | 0.779 | 0.407 | 0.978 | 0.448 | 0.064 | 0.552 |
| | Attention | 0.557 | 0.557 | [0.543, 0.569] | 0.878 | 0.176 | 0.968 | 0.727 | 0.058 | 0.273 |

*correlation between baseline and follow-up #1 corrected for participant age, timepoint and their interaction. ICC = Interclass correlation coefficient; CBCL = Child Behavior Checklist; S = specific factors; GP = general psychopathology; DP = dysregulation profile; omega reliability index; xH = omega-hierarchical; FD = factor determinacy; ECV = explained common variance; SS = proportion of common variance of the items in each factor that is due to that factor; SG = ECV proportion of common variance of the items in each specific factor that is due to the specific factor; GS = proportion of common variance of the items in each specific factor that is due to the general factor

#### Factor and summary score reliability

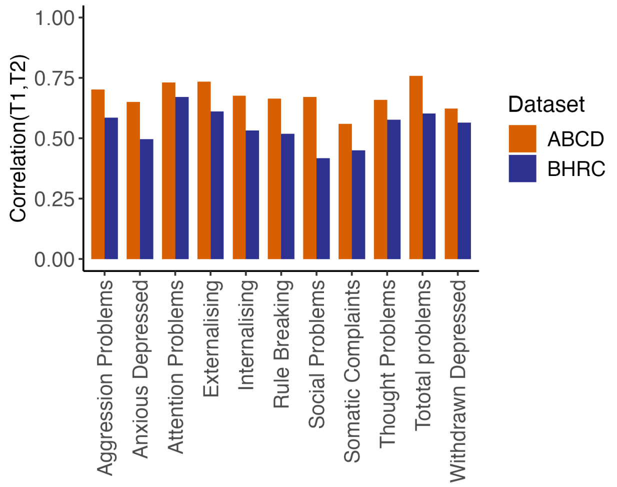


Supplementary Figure 3. Summary Score Reliability: Test-retest Correlation

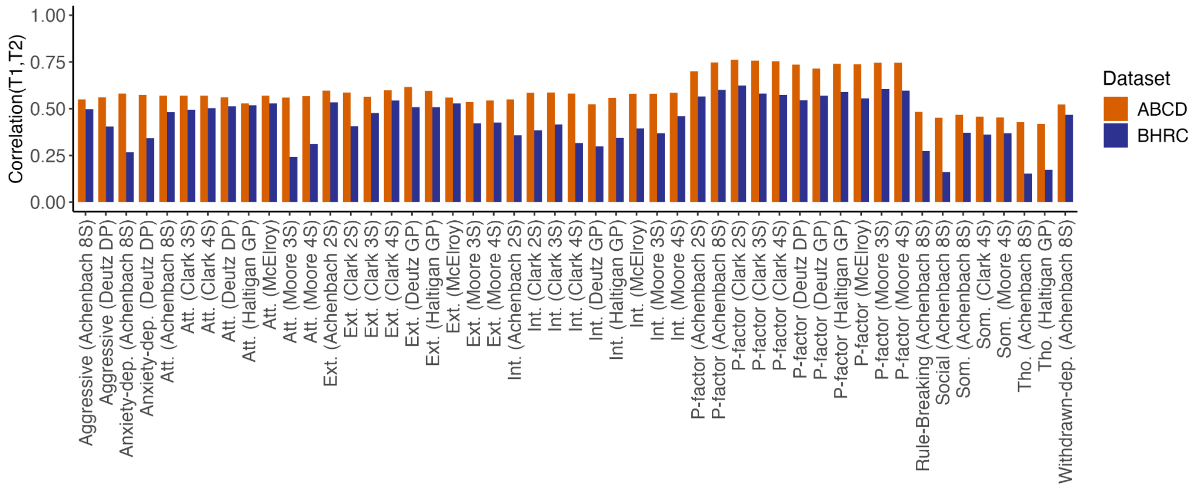


Supplementary Figure 4. Bifactor Reliability: Test-retest Correlation

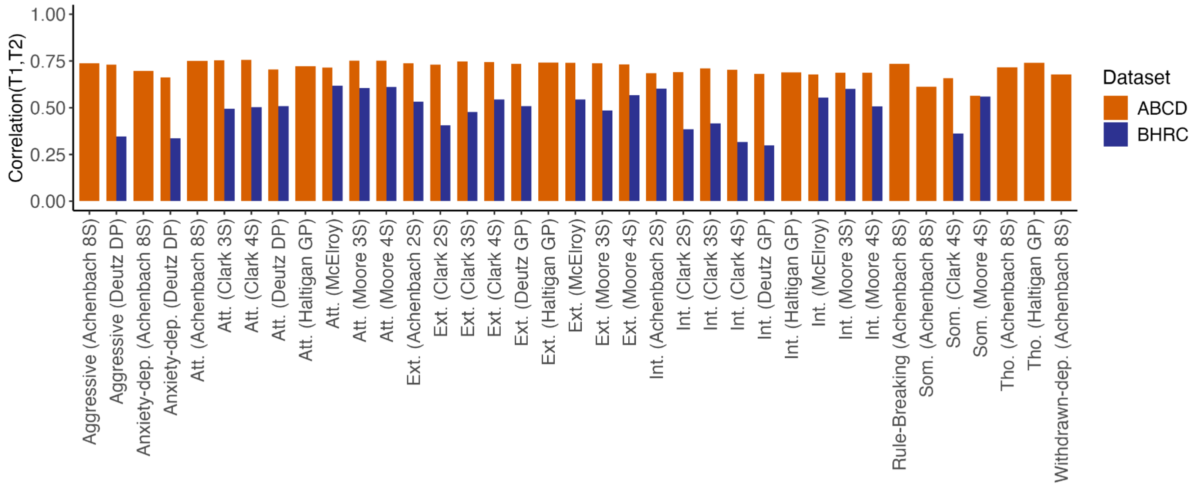


Supplementary Figure 5. Correlated Factor Reliability: Test-retest Correlation

#### Supplement to the prediction of factor and summary scores from functional connectivity and cortical thickness

**Supplementary Table 5**

*Bifactor score prediction accuracy and reliability*

| Factor (model) | Categorisation | Test-retest correlation | ICC | Functional connectivity | | Cortical thickness | |
|---|---|---|---|---|---|---|---|
| | | | | $R^2$ | Pearson r | $R^2$ | Pearson r |
| P-factor (Achenbach 2S) | P-Factor | 0.700 | 0.700 | 0.008 | 0.092* | 0.001 | 0.052* |
| Int. (Achenbach 2S) | Internalising | 0.550 | 0.550 | -0.004 | 0.048* | -0.004 | 0.011 |
| Ext. (Achenbach 2S) | Externalising | 0.596 | 0.596 | 0.019 | 0.147* | 0.002 | 0.060* |
| P-factor (Achenbach 8S) | P-Factor | 0.748 | 0.748 | 0.013 | 0.120* | 0.003 | 0.073* |
| Anxiety-dep. (Achenbach 8S) | Other | 0.580 | 0.580 | 0.018 | 0.137* | -0.0002 | 0.049* |
| Withdrawn-dep. (Achenbach 8S) | Other | 0.523 | 0.524 | 0.003 | 0.077* | -0.002 | 0.060* |
| Som. (Achenbach 8S) | Other | 0.467 | 0.467 | 0.001 | 0.042 | -0.0003 | 0.001 |
| Rule-Breaking (Achenbach 8S) | Other | 0.483 | 0.482 | 0.010 | 0.106* | 0.001 | 0.062* |
| Aggressive (Achenbach 8S) | Other | 0.550 | 0.550 | 0.009 | 0.097* | 0.0001 | 0.019 |
| Social (Achenbach 8S) | Other | 0.452 | 0.452 | 0.003 | 0.065* | 0.002 | 0.054* |
| Tho. (Achenbach 8S) | Other | 0.429 | 0.428 | -0.006 | 0.011 | -0.002 | 0.008 |
| Att. (Achenbach 8S) | Attention | 0.570 | 0.570 | 0.009 | 0.096* | -0.002 | 0.024 |
| P-factor (Moore 3S) | P-Factor | 0.747 | 0.747 | 0.011 | 0.112* | 0.002 | 0.063* |
| Int. (Moore 3S) | Internalising | 0.579 | 0.580 | 0.006 | 0.087* | 0.00002 | 0.035 |
| Ext. (Moore 3S) | Externalising | 0.537 | 0.537 | 0.011 | 0.107* | 0.001 | 0.035 |
| Att. (Moore 3S) | Attention | 0.560 | 0.560 | 0.010 | 0.102* | -0.002 | 0.038 |
| P-factor (Moore 4S) | P-Factor | 0.746 | 0.746 | 0.012 | 0.114* | 0.002 | 0.062* |
| Int. (Moore 4S) | Internalising | 0.585 | 0.586 | -0.012 | 0.109* | -0.026 | 0.012 |

| | | | | | | | |
|---|---|---|---|---|---|---|---|
| Som. (Moore 4S) | Other | 0.453 | 0.453 | 0.001 | 0.047 | -0.001 | -0.005 |
| Ext. (Moore 4S) | Externalising | 0.545 | 0.544 | 0.009 | 0.097* | 0.001 | 0.033 |
| Att. (Moore 4S) | Attention | 0.567 | 0.567 | 0.010 | 0.103* | -0.001 | 0.042 |
| P-factor (McElroy) | P-Factor | 0.738 | 0.738 | 0.009 | 0.105* | 0.002 | 0.069* |
| Int. (McElroy) | Internalising | 0.579 | 0.579 | 0.008 | 0.092* | 0.001 | 0.034 |
| Ext. (McElroy) | Externalising | 0.560 | 0.559 | 0.013 | 0.126* | -0.001 | 0.047* |
| Att. (McElroy) | Attention | 0.569 | 0.569 | 0.008 | 0.095* | -0.001 | 0.042 |
| P-factor (Deutz GP) | P-Factor | 0.715 | 0.715 | 0.009 | 0.096* | 0.002 | 0.050* |
| Int. (Deutz GP) | Internalising | 0.525 | 0.525 | 0.003 | 0.079* | -0.002 | 0.033 |
| Ext. (Deutz GP) | Externalising | 0.617 | 0.617 | 0.017 | 0.138* | -0.001 | 0.043 |
| P-factor (Deutz DP) | P-Factor | 0.735 | 0.735 | 0.012 | 0.110* | 0.002 | 0.062* |
| Anxiety-dep. (Deutz DP) | Internalising | 0.573 | 0.573 | 0.013 | 0.118* | -0.002 | 0.028 |
| Aggressive (Deutz DP) | Externalising | 0.563 | 0.562 | 0.009 | 0.098* | 0.0003 | 0.021 |
| Att. (Deutz DP) | Attention | 0.562 | 0.562 | 0.010 | 0.100* | 0.001 | 0.053* |
| P-factor (Haltigan GP) | P-Factor | 0.739 | 0.740 | 0.011 | 0.107* | 0.002 | 0.060* |
| Int. (Haltigan GP) | Internalising | 0.558 | 0.558 | 0.002 | 0.068* | -0.001 | 0.023 |
| Ext. (Haltigan GP) | Externalising | 0.595 | 0.594 | 0.016 | 0.139* | -0.001 | 0.038 |
| Tho. (Haltigan GP) | Other | 0.418 | 0.416 | -0.016 | -0.030 | -0.013 | 0.003 |
| Att. (Haltigan GP) | Attention | 0.528 | 0.528 | 0.007 | 0.095* | -0.002 | 0.040 |
| P-factor (Clark 2S) | P-Factor | 0.762 | 0.762 | 0.017 | 0.139* | 0.004 | 0.078* |
| Int. (Clark 2S) | Internalising | 0.585 | 0.586 | 0.010 | 0.103* | 0.001 | 0.046* |
| Ext. (Clark 2S) | Externalising | 0.588 | 0.587 | 0.013 | 0.118* | -0.0002 | 0.023 |
| P-factor (Clark 3S) | P-Factor | 0.758 | 0.758 | 0.015 | 0.125* | 0.004 | 0.068* |
| Int. (Clark 3S) | Internalising | 0.587 | 0.588 | -0.013 | 0.097* | -0.020 | 0.040 |
| Ext. (Clark 3S) | Externalising | 0.563 | 0.562 | 0.012 | 0.112* | 0.001 | 0.038 |
| Att. (Clark 3S) | Attention | 0.569 | 0.569 | 0.006 | 0.102* | -0.006 | 0.029 |
| P-factor (Clark 4S) | P-Factor | 0.753 | 0.753 | 0.013 | 0.120* | 0.003 | 0.069* |
| Int. (Clark 4S) | Internalising | 0.582 | 0.582 | 0.018 | 0.136* | -0.003 | 0.026 |
| Som. (Clark 4S) | Other | 0.458 | 0.458 | 0.001 | 0.046 | -0.0004 | 0.002 |
| Ext. (Clark 4S) | Externalising | 0.599 | 0.599 | 0.011 | 0.108* | 0.001 | 0.034 |
| Att. (Clark 4S) | Attention | 0.570 | 0.570 | 0.009 | 0.094* | -0.002 | 0.030 |

P-Factor = the general factor of psychopathology; GP = general psychopathology model; DP = dysregulation profile model; * designates permutation-based significant predictions at p<0.001

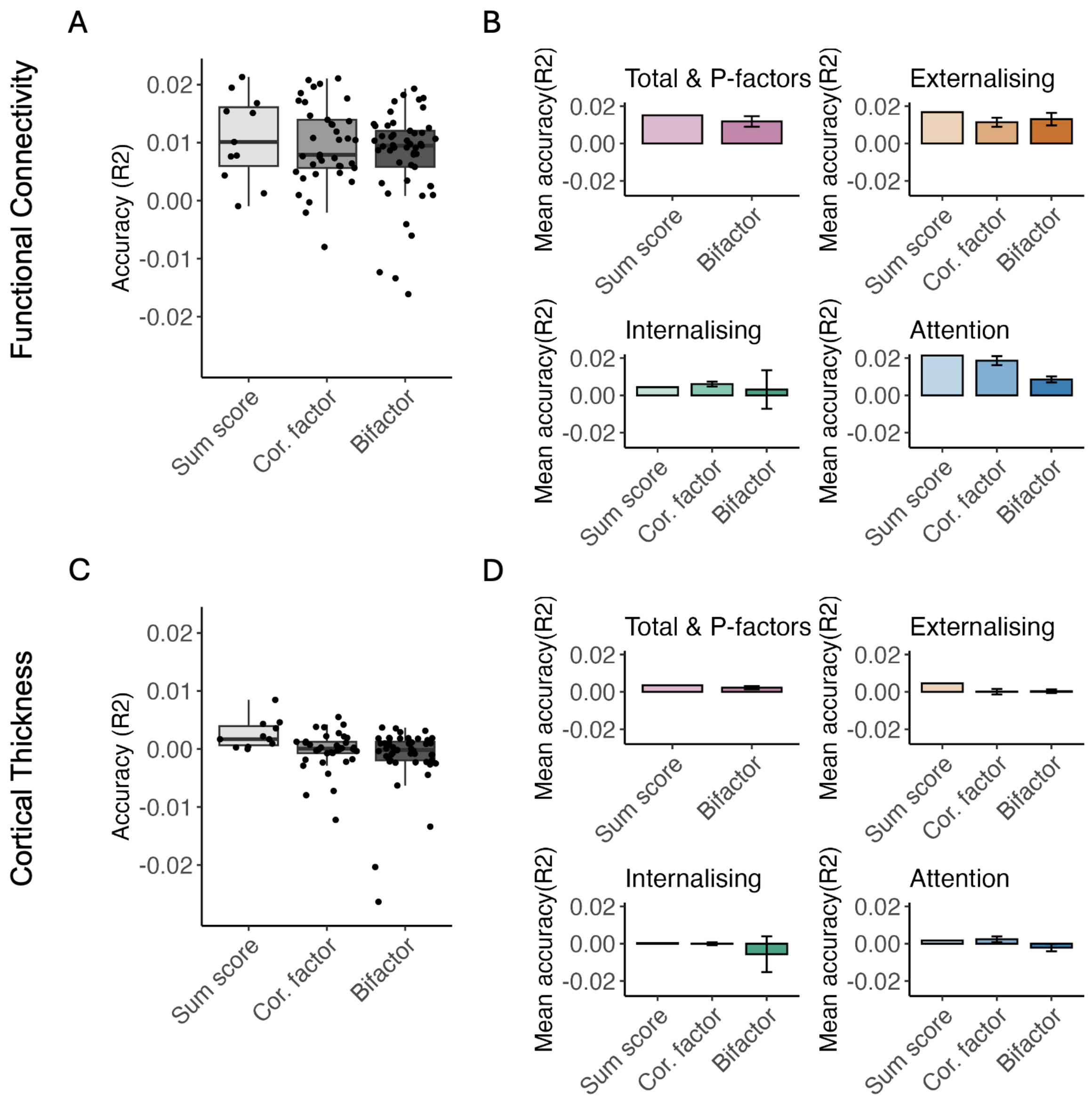


Supplementary Figure 6. Prediction accuracy of CBCL summary and factor scores: Coefficient of Determination

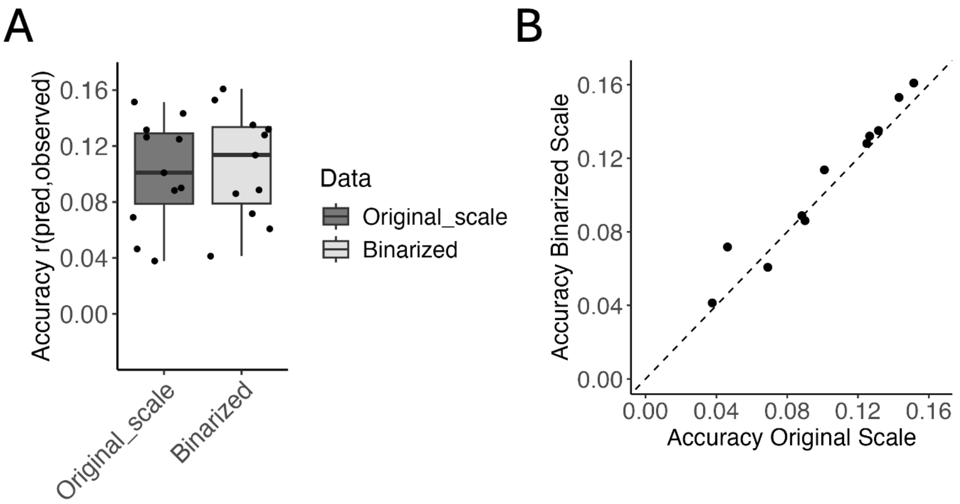


Supplementary Figure 7. Comparison of summary scores calculated using rescored (i.e, binarised) and original CBCL items. Panel A shows the distribution of all prediction accuracies for both calculated summary scores. Panel B shows the prediction accuracy for both calculated summary scores. The dashed line represents the identity line.

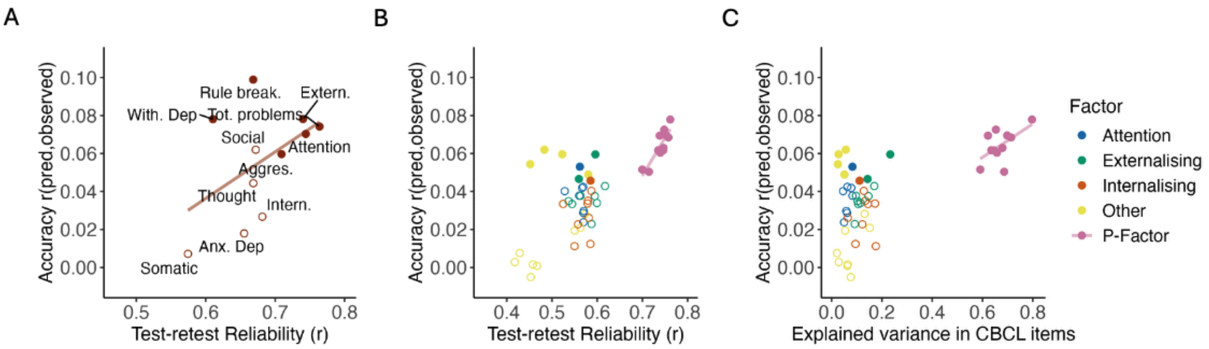


Supplementary Figure 8. Impact of psychometrics on the prediction accuracy of bifactor scores by cortical thickness.

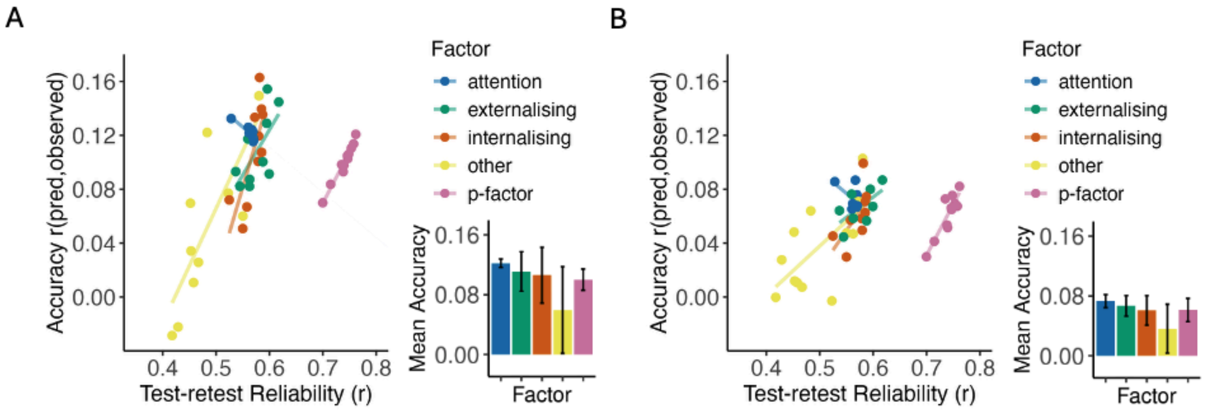


Supplementary Figure 9. Control analyses for the prediction of bifactor scores from functional connectivity. Panel A shows the prediction of factor scores calculated on follow-up 2 CBCL data in the ABCD dataset. Panel B details prediction accuracy using XGBoost in place of ridge regression.

The distribution of prediction accuracies using XGBoost closely mirrored those obtained with ridge regression, albeit with lower performance, indicating that within each factor group (i.e., P, externalising, internalising, attention) increasing reliability generally resulted in higher prediction accuracy; however, between factor groups, P-factors and most specific factors could be predicted with similar accuracy across machine learning models (Supplementary Figure 8A). Additionally, this pattern was consistent across longitudinal timepoints when using factors from follow-up 1 CBCL data (Supplementary Figure 8B). Taken together, these findings suggest that although general factors were more psychometrically robust, this alone did not necessarily enhance their association with neuroimaging features.

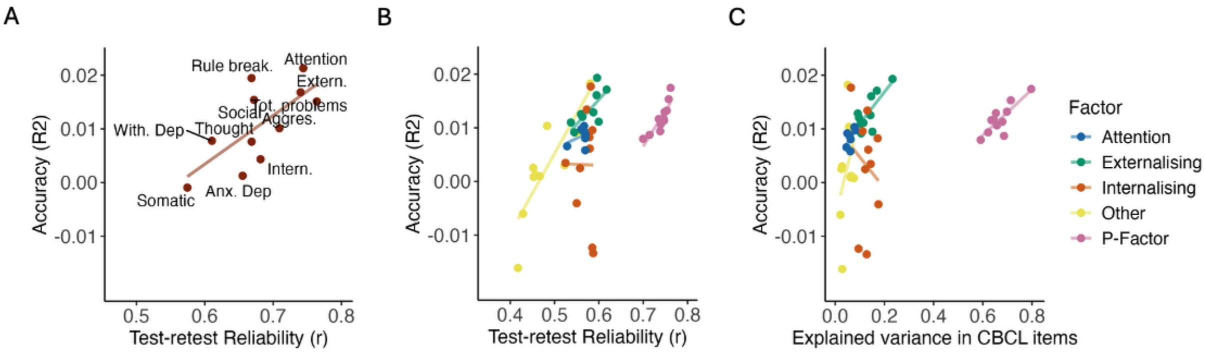


Supplementary Figure 10. Impact of psychometrics on the prediction accuracy of bifactor scores by functional connectivity – Coefficient of determination

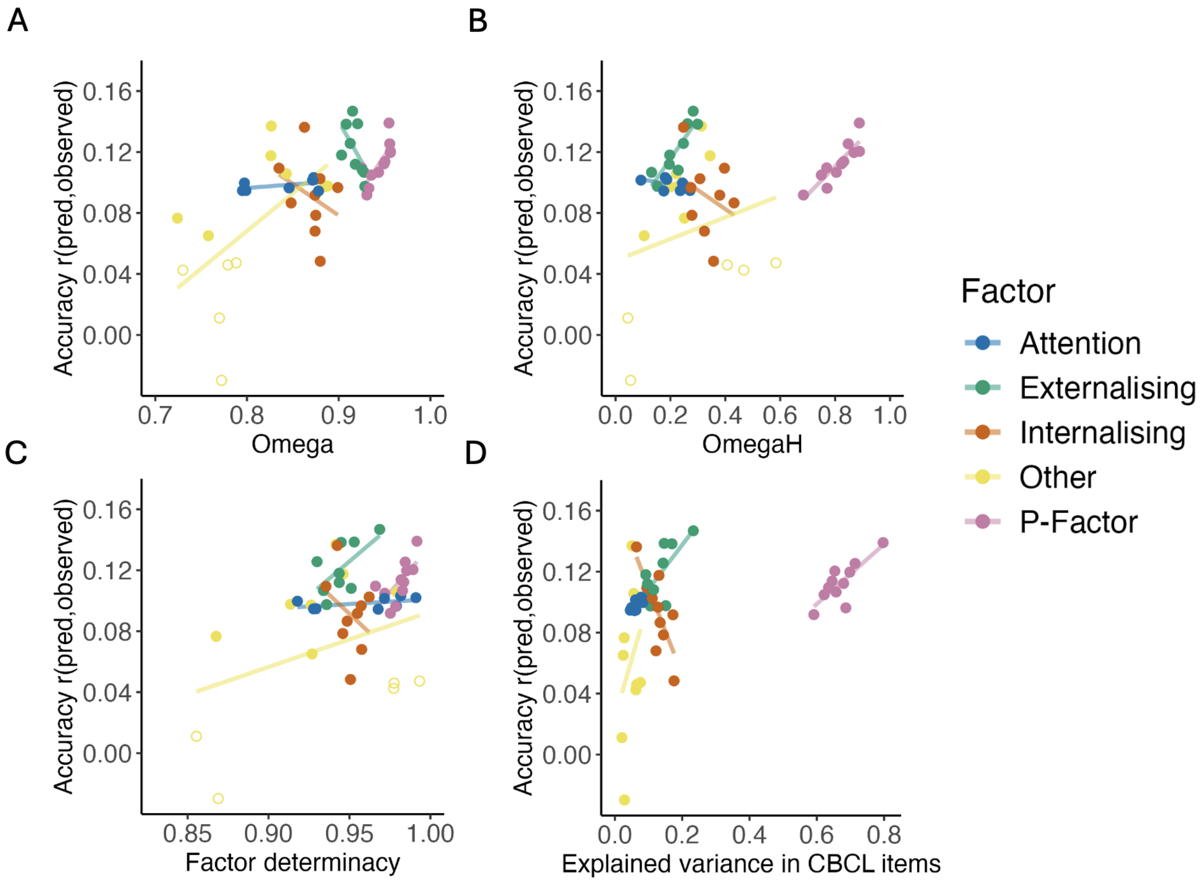


Supplementary Figure 11. Impact of internal consistency metrics on the prediction accuracy of bifactor scores by functional connectivity.

Replicating observations from other datasets (Hoffmann et al., 2022) and evident from Supplementary Figure 10, internal consistency across most P-factors in the ABCD was high ($Omega_{mean}$ = 0.94, Omega = 0.93 - 0.96; $omegaH_{mean}$ = 0.81, omegaH = 0.68 - 0.88), while specific factors showed low to acceptable internal consistency (Omega: mean = 0.85, 0.72 - 0.93; omegaH: mean = 0.26, 0.05 - 0.58). Notably, all three metrics displayed a substantial correlation with test-retest reliability ($\rho_{Omega}$ = 0.77, p < 0.001; $\rho_{omegaH}$ = 0.64, p < 0.001; $\rho_{FD}$ = 0.50, p < 0.001).

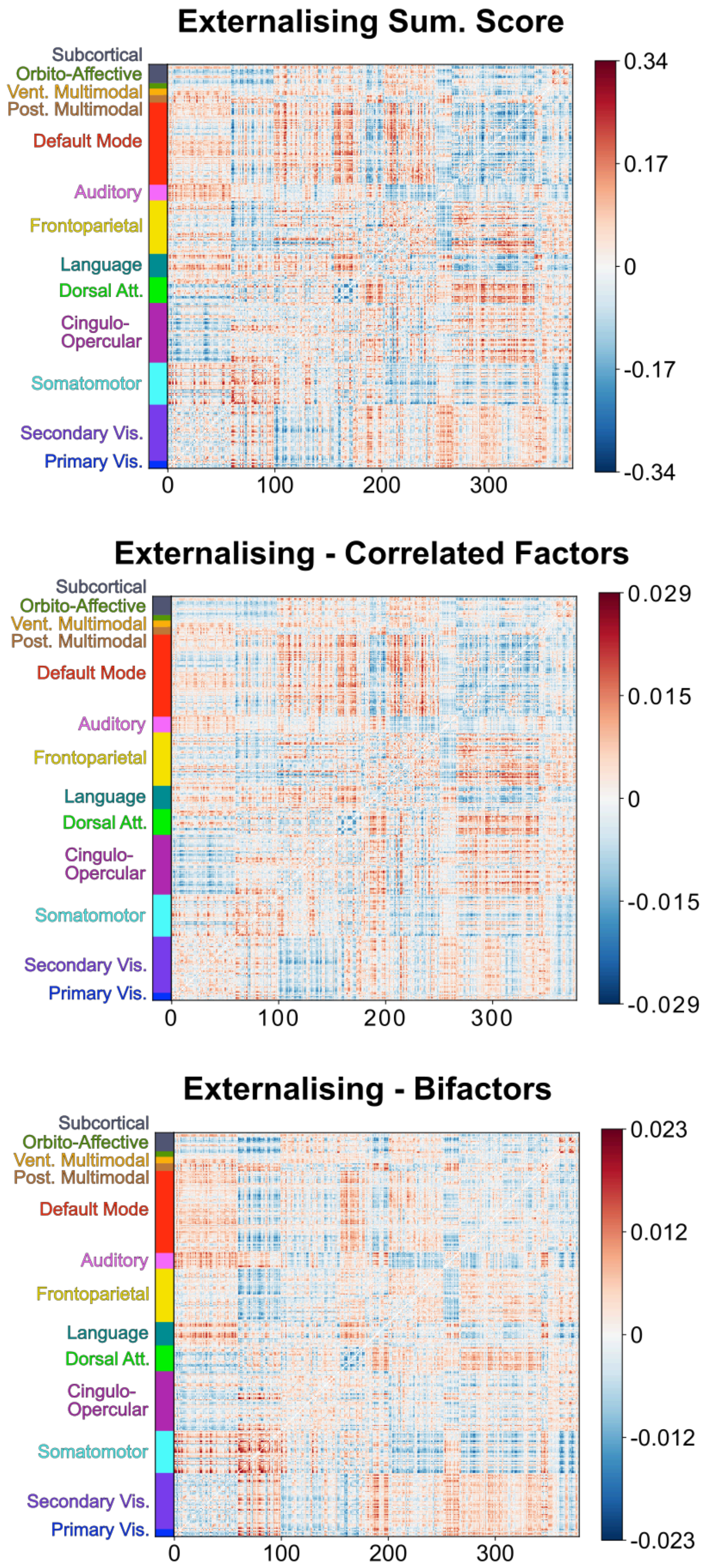


Supplementary Figure 12. Haufe-transformed feature importance weights of edges between all cortical parcels: Comparison of externalising factor and summary score models. Positive or negative feature weight for an edge indicates that higher connectivity for that edge was associated with predicting higher or lower behavioural value, respectively. Ordered using the functional network definition by Ji et al. (2019). Note, weights for bifactor and correlated factor models are averaged across bifactor and correlated factor model solutions, respectively.

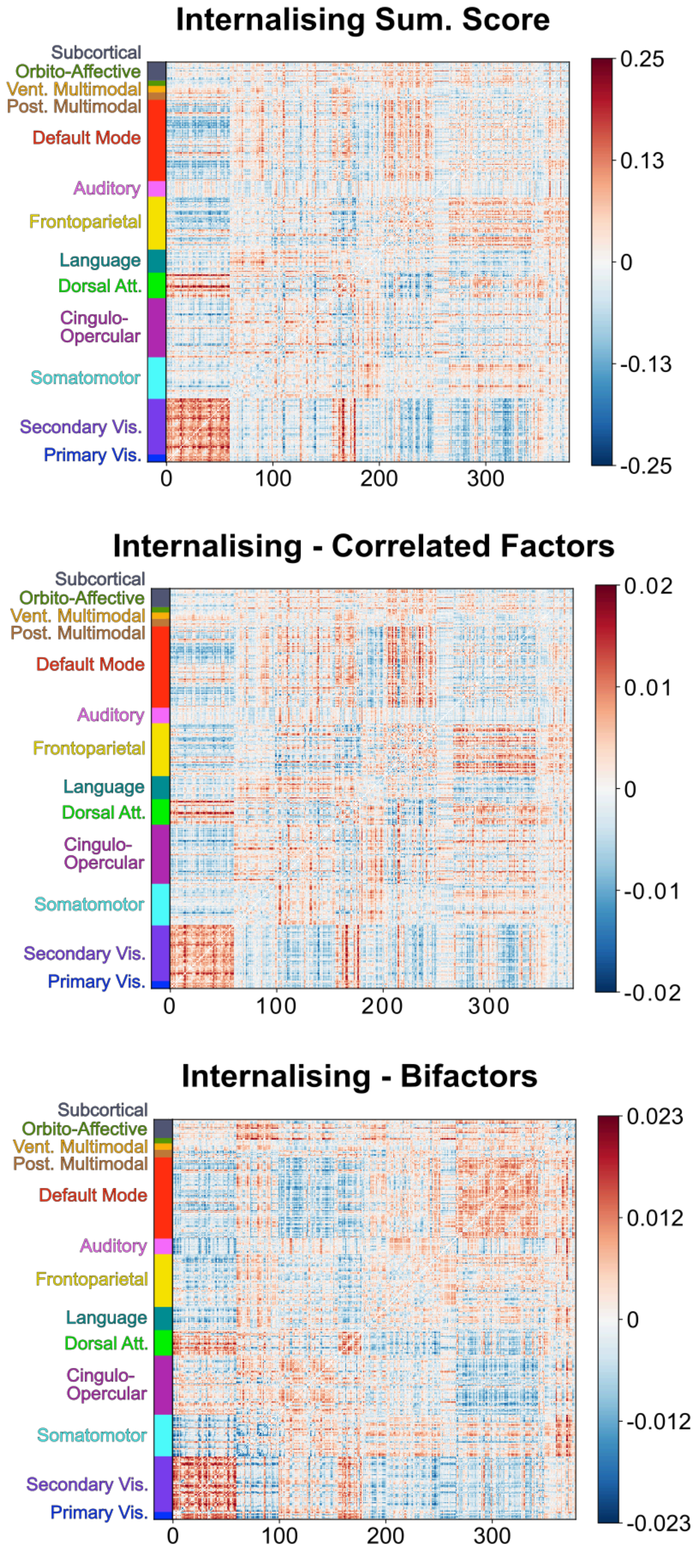


Supplementary Figure 13. Haufe-transformed feature importance weights of edges between all cortical parcels: Comparison of internalising factor and summary score models. Positive or negative feature weight for an edge indicates that higher connectivity for that edge was associated with predicting higher or lower behavioural value, respectively. Ordered using the functional network definition by Ji et al. (2019). Note that weights for bifactor and correlated factor models are averaged across bifactor and correlated factor model solutions, respectively.

#### Factor correlations

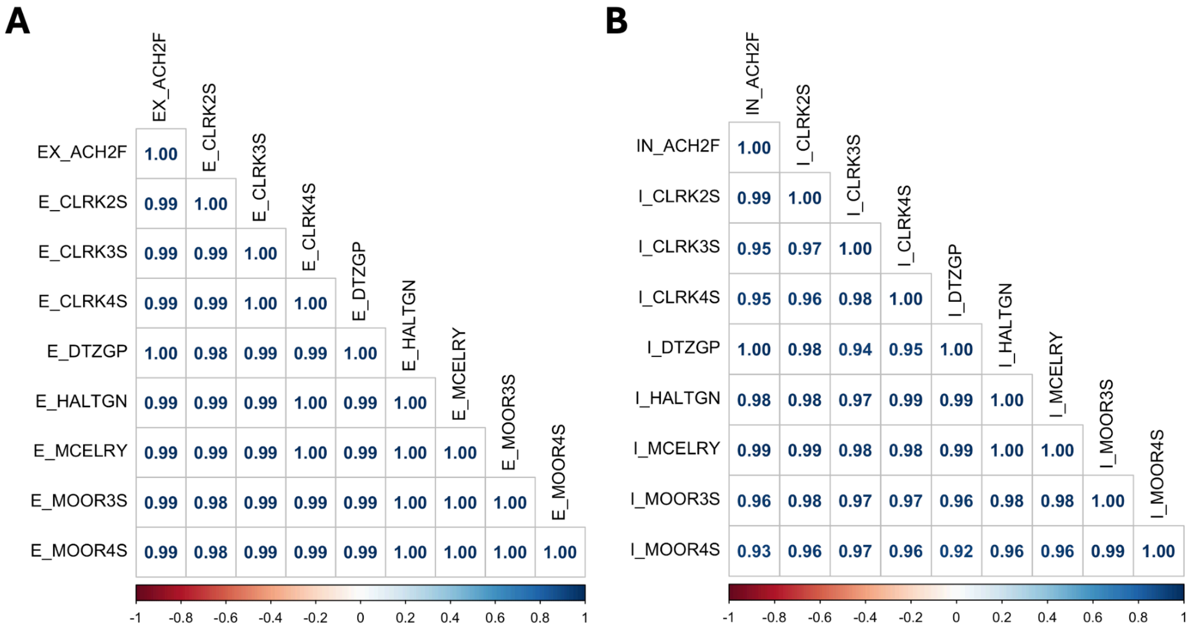


Supplementary Figure 14. Consistency in feature weights across correlated factor model solutions

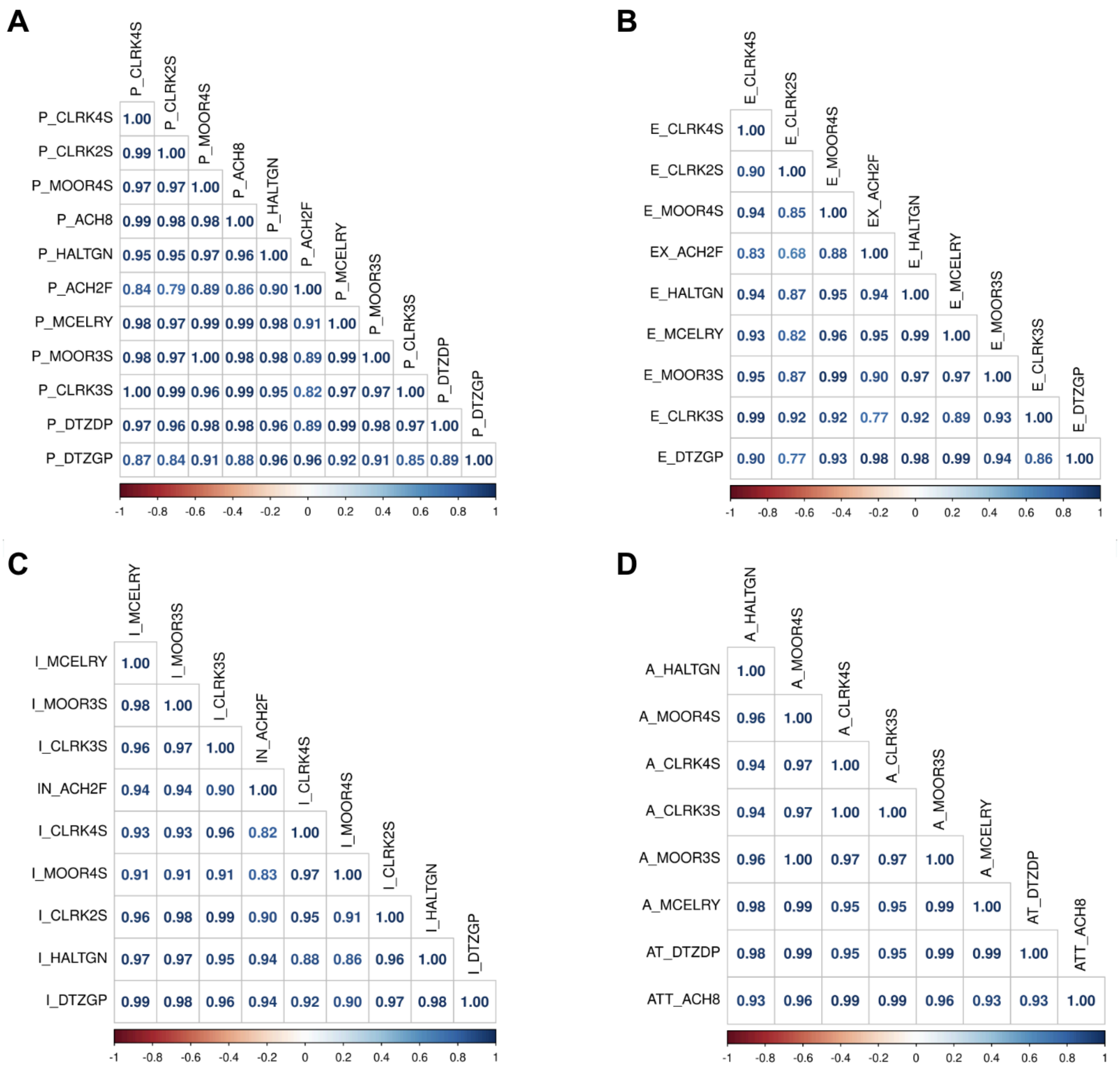


Supplementary Figure 15. Consistency in feature weights across bifactor model solutions

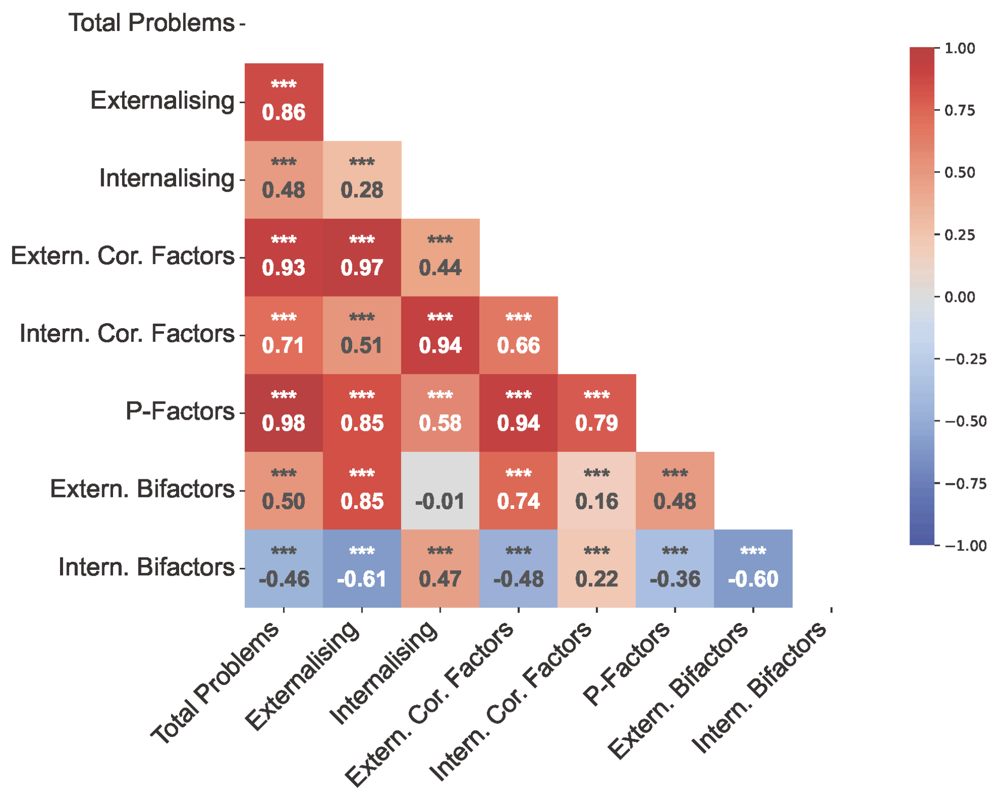


Supplementary Figure 16. Correlation between informative features for the prediction of all factors and summary scores.

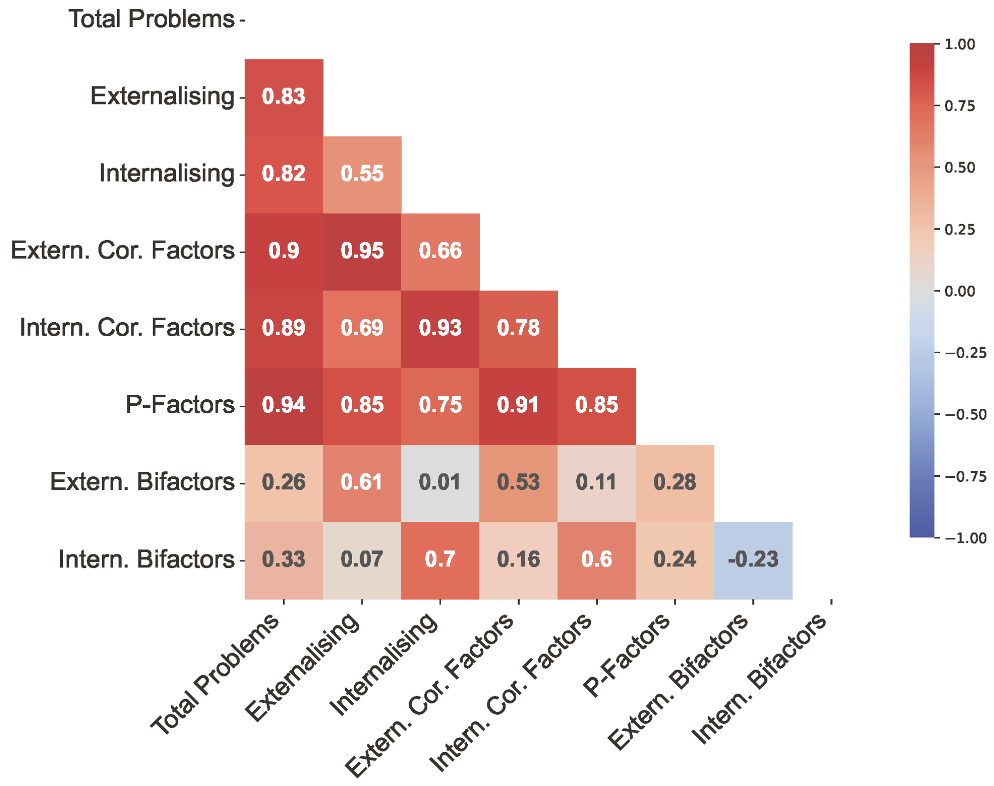


Supplementary Figure 17. Correlation between all factors and summary scores.

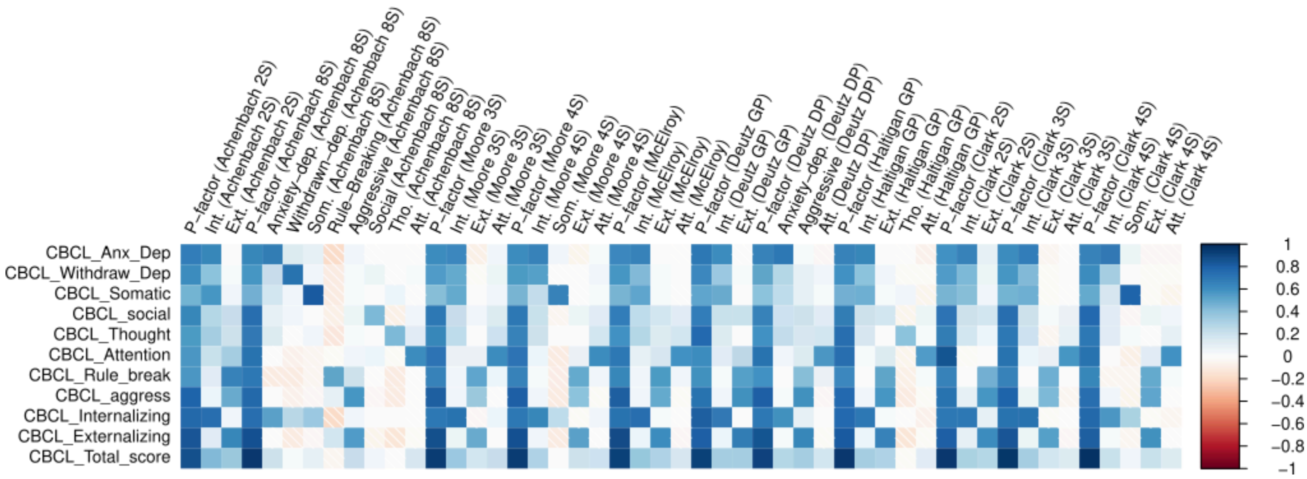


Supplementary Figure 18. Correlations between mean bifactor scores and all summary scores

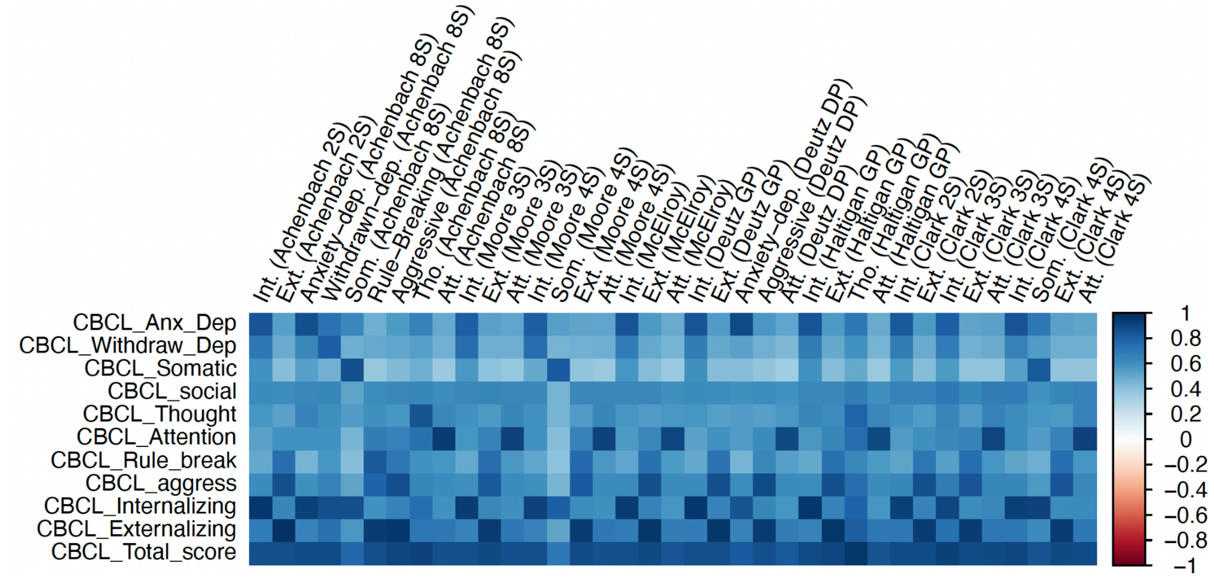


Supplementary Figure 19. Correlations between mean correlated factor scores and all summary scores

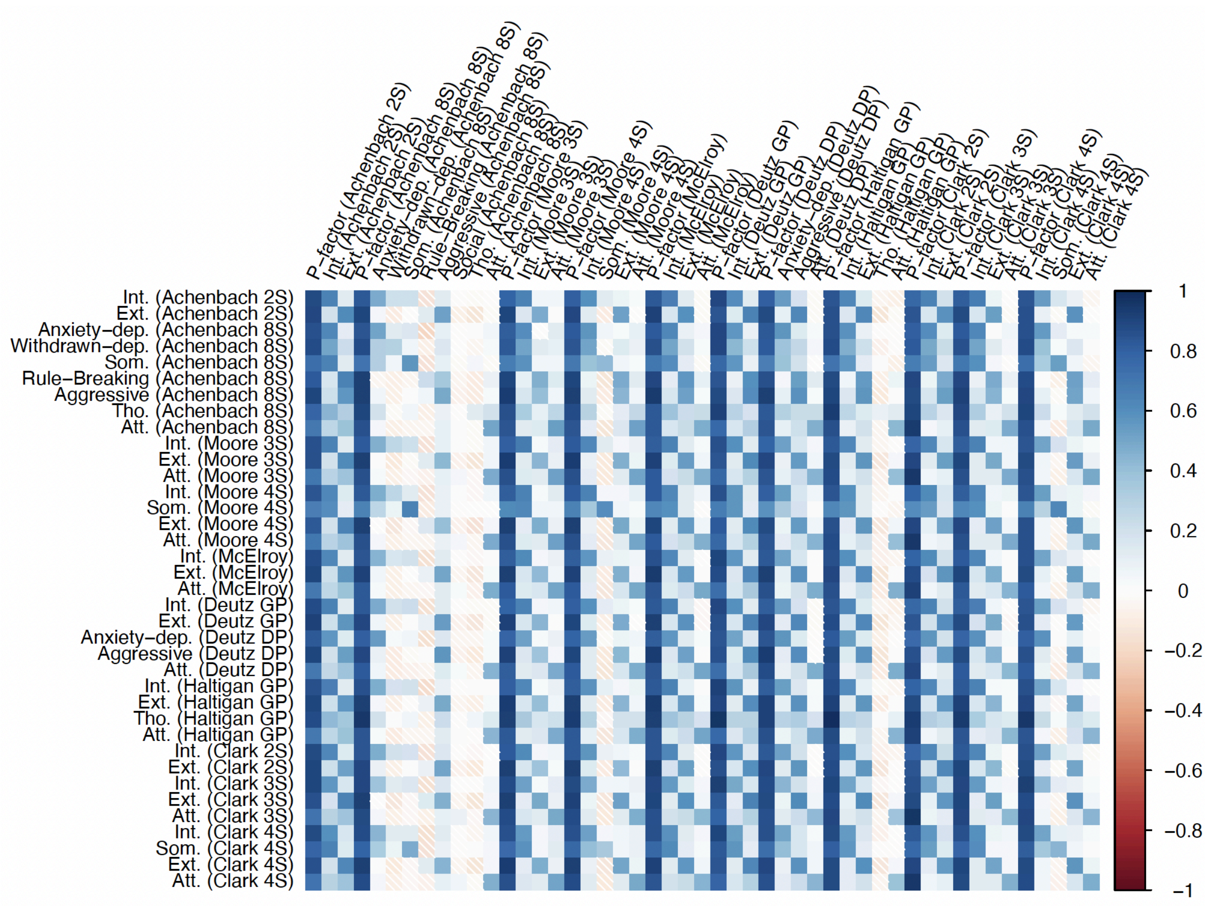


Supplementary Figure 20. Correlations between mean bifactor scores and mean correlated factor scores.
